# Phase- and State-Dependent Modulation of Breathing Pattern by preBötzinger Complex Somatostatin Expressing Neurons

**DOI:** 10.1101/2021.04.12.439520

**Authors:** Raquel P. de Sousa Abreu, Evgeny Bondarenko, Jack L. Feldman

**Affiliations:** Department of Neurobiology, David Geffen School of Medicine, University of California at Los Angeles, Los Angeles, CA, 90095

**Keywords:** preBötC, breathing pattern, optogenetics, somatostatin

## Abstract

As neuronal subtypes are increasingly categorized, delineating their functional role is paramount. The preBötzinger Complex (preBötC) subpopulation expressing the neuropeptide somatostatin (SST) is classified as mostly excitatory, inspiratory-modulated and not rhythmogenic. We further characterized their phenotypic identity; 87% were glutamatergic and the balance were glycinergic and/or GABAergic. We then used optogenetics to investigate their modulatory role in both anesthetized and freely moving mice. In anesthetized mice, short photostimulation (100 ms) of preBötC SST^+^ neurons modulated breathing-related variables in a combinatory phase- and state-dependent manner; changes in inspiratory duration, inspiratory peak amplitude (Amp), and phase were different at higher (≥2.5 Hz) vs. lower (<2.5 Hz) breathing frequency. Moreover, we observed a biphasic effect of photostimulation during expiration that is probabilistic, i.e., photostimulation given at the same phase in consecutive cycles can evoke opposite responses (lengthening vs. shortening of the phase). This unexpected probabilistic state- and phase-dependent responses to photostimulation exposed properties of the preBötC that were not predicted and cannot be readily accounted for in current models of preBötC pattern generation. In freely moving mice, prolonged photostimulation decreased *f* in normoxia, hypoxia, or hypercapnia, and increased Amp and produced a phase advance, which was similar to the results in anesthetized mice when *f*≥2.5 Hz. We conclude that preBötC SST^+^ neurons are a key mediator of the extraordinary and essential lability of breathing pattern.

**Key points summary:**

- We transfected preBötzinger Complex (preBötC) somatostatin-expressing (SST^+^) neurons, which modulate respiratory pattern, but are not rhythmogenic, with channelrhodopsin to investigate phase- and state-dependent modulation of breathing pattern in anesthetized and freely behaving mice in normoxia, hypoxia, and hypercapnia.
- In anesthetized mice, photostimulation of preBötC SST^+^ neurons during inspiration increased inspiratory duration and amplitude regardless of baseline breathing frequency, *f*.
- In anesthetized mice with low *f* (<2.5 Hz), photostimulation of preBötC SST^+^ neurons during expiration evoked either phase advance or phase delay, whereas in anesthetized mice with high *f* (≥2.5 Hz) and in freely behaving mice in normoxia, hypoxia, or hypercapnia, photostimulation always evoked phase advance.
- Phase- and state-dependency is a function of overall breathing network excitability.
- The *f*-dependent probabilistic modulation of breathing pattern by preBötC SST^+^ neurons was unexpected, requiring reconsideration of current models of preBötC function, which neither predict nor can readily account for such responses.

## Introduction

The preBötC, the kernel for the breathing central pattern generator (bCPG) in mammals, q.v., (Smith *et al*., 1991; Del Negro, Funk and Feldman, 2018), is a neural microcircuit with numerous subpopulations of glutamatergic excitatory and glycinergic and GABAergic inhibitory neurons. These subpopulations are demarked by the expression of various molecular markers, e.g., SST, reelin, SST receptor, Dbx1, neurokinin 1 receptor, neuromedin B receptor, gastrin-releasing peptide receptor, μ-opioid receptor (Gray *et al*., 1999, 2001; Liu, Ju and Wong-Riley, 2001; Stornetta *et al*., 2003; Stornetta, 2008; Tan *et al*., 2010; Feldman and Kam, 2015; Li *et al*., 2016). Here we focus on a subset of preBötC glutamatergic neurons that expresses SST (SST^+^ neurons) that are presumably non-rhythmogenic (Cui *et al*., 2016; Ashhad and Feldman, 2020), but convey excitatory preBötC output to multiple sites, including to premotor inspiratory neurons (Cui *et al*., 2016; Yang and Feldman, 2018; Yang *et al*., 2020). Using optogenetic excitation of preBötC SST^+^ neurons in anesthetized and freely moving mice, we demonstrate that these neurons not only transmit signals from the preBötC to efferent targets to ultimately produce inspiratory muscle activity, but also modulate its activity.

Since various parameters of breathing pattern, e.g., cycle duration, frequency, tidal volume, can vary rapidly (breath by breath) and considerably during normal rodent motor behavior (Kabir *et al*., 2010), e.g., due to epochs of grooming, chewing, and walking, our first approach was to study anesthetized but otherwise intact mice to reduce these confounds. Taking advantage of the temporal precision of optogenetic photostimulation that can illuminate dynamic properties of the bCPG (Sherman *et al*., 2015; Cui *et al*., 2016), we investigated the effects on breathing-related variables of briefly (100 ms) exciting preBötC SST^+^ neurons at different times during the breathing cycle. Photostimulation during inspiration increased inspiratory duration (T_I_) and/or amplitude (Amp) while photostimulation during expiration shortened onset to next inspiration, i.e., produced a breathing cycle phase advance. These effects were dependent on baseline breathing frequency, *f*, with a marked difference below or above 2.5 Hz. For instance, photostimulation during expiration had a bimodal effect on the phase, i.e., advance or delay, when *f*<2.5 Hz but a unimodal effect, i.e., phase advance, when *f*≥2.5 Hz. Thus, SST-mediated modulation of breathing was both phase- and state-dependent.

We extended these studies to unanesthetized freely moving adult mice, breathing normoxic (room air), hypoxic (8% O_2_) and hypercapnic (5% CO_2_) gas mixtures. In normoxia, hypoxia, and hypercapnia, photostimulation had similar effects on *f*, the coefficient of variation of breathing cycle duration, and Amp. In general, during photostimulation, *f* and variability significantly decreased. These results were consistent with results in anesthetized mice when *f*≥2.5 Hz.

Photostimulation of preBötC SST^+^ neurons changed breathing pattern with a strong dependence on phase timing of the stimulus and behavioral state. Given this dependence, we suggest that preBötC SST^+^ neurons can play a powerful modulatory role in both rapid and slow breathing responses, such as during startle, exercise, stress, disease, and changes of blood gases or altitude, that contribute to the exceptional lability of breathing pattern.

## Materials and Methods

### Animal and virus usage

A SST-IRES-cre mouse line (strain Sst^tm2.1(cre)Zjh^/J, # 013044, The Jackson Laboratory; (Taniguchi *et al*., 2011)) was maintained at the Division of Laboratory Animal Medicine of the University of California, Los Angeles (UCLA). Mice were housed on a 12/12 h light/dark cycle in a temperature and humidity controlled room (70-76°F, 30-70%, respectively) and with *ad libitum* access to food and water. Animal use and experimental protocols were approved by the University of California at Los Angeles Animal Research Committee (#A3196-01, #1994-159-91). All efforts were made to minimize animal suffering and discomfort and to reduce the number of animals used. Prior to surgery, animals were housed in groups of variable size up to five mice per cage; after surgery, animals were housed individually as required by our study protocols.

To selectively transfect preBötC SST^+^ neurons with Channelrhodopsin (ChR2), we used the Cre-inducible recombinant viral vector (AAV2/1-EF1α-DIO-hChR2(H134R)-eYFP-WPRE-hGH; University of Pennsylvania Vector Core), which carries a floxed and enhanced version of the photosensitive cationic channel ChR2 fused with the enhanced Yellow Fluorescent protein (eYFP). Since the vector injected is flanked by double lox sites (loxP and lox2722) with an inverted reading frame that is Cre-dependent, only in the presence of Cre recombination and inversion into the sense direction occurs and functional expression is acquired.

### Surgical procedures

Pre-operatively, SST-cre male mice (10 weeks old) received a dose of buprenorphine (0.05-0.1 mg/Kg). Animals were then deeply anesthetized with isoflurane (4%) and placed in a stereotaxic apparatus (David Kopf Instruments, Tujunga, CA, USA) using Bregma and Lambda as skull landmarks. Anesthesia, ≈2% isoflurane, was maintained throughout the surgical procedure via a nose cone. Animals were placed on a heating pad and temperature of the animal was monitored by a rectal probe that was connected to a rodent temperature controller (TCAT 2DF, Physitemp Instruments, Inc.) to maintain the animal at 37^°^C. Craniotomy was performed to stereotaxically inject up to 50 nL of the cre-dependent AAV2 virus uni- or bilaterally into the preBötC with a glass micropipette and an air pressure injector system (Picospritzer II, Parker Hannafin), similarly to established methods (Sherman *et al*., 2015; Cui *et al*., 2016). The micropipette was left in place for 10 to 15 minutes after each injection to allow viral diffusion and to minimize backflow of the virus up the pipette track. The wound was closed with non-absorbable sutures. Post-operatively, mice were housed individually and received buprenorphine (0.05-0.1 mg/Kg) at 12 hours intervals for 48 hours and sulfamethoxadozole and trimethoprim (TMS) in drinking water during the following week. One to two weeks after the virus injection, a second surgical procedure was performed as described above. Fiber-optic cannulas with a 1.25 mm stainless steel ferrule (200μm, Doric Lenses or ThorLabs) were stereotaxically placed dorsally to the preBötC and fixed to the skull with dental cement (C&B-METABOND®; Parkell inc.). The wound was sealed by applying a layer of dental acrylic (Ortho-Jet powder, Ortho-Jet liquid, Lang Dental).

### RNA analysis (RNAscope)

10-14 week old SST-Cre mice were euthanised with isoflurane overdose, their brainstems were rapidly removed and flash frozen in dry ice. 20 µm sagittal sections of fresh frozen brainstems were cut on a cryostat (CryoStar NX70, Thermo Scientific), mounted on SuperFrost Plus (Fisher Scientific) slides and stored at −80 °Cfor at least 1 day before processing. We followed manufacturer’s protocol for fluorescent *in situ* hybridization (RNAscope version 1, Advanced Cell Diagnostics). Briefly, tissue samples were postfixed in 10% neutral buffered paraformaldehyde, washed, and dehydrated in sequential concentrations of ethanol (50, 70, and 100%). Samples were treated with protease IV and incubated for 2 hours at 40 C in HybEZ^TM^ Hybridization Oven (Advanced Cell Diagnostics) in the presence of target probes. We used combinations of following probes for hybridization: Mm-ChAT, *Mm-Sst*, *Mm-Slc17a6 (VGluT2), Mm-Slc6a5 (GlyT2), Mm-Gad1* and *Cre*. Probe for Mm-ChAT was used in all probe combinations to mark the location of nucleus ambiguus, which is necessary for determining the location of preBötC. After a 4-step amplification process, samples were counterstained with DAPI and coverslipped with ProLong Gold (Invitrogen) used as a mounting agent. Images were acquired on a confocal laser scanning microscope (LSM710 META, Zeiss). High-resolution z-stack confocal images were taken at 1 µm intervals and then merged using ImageJ software.

Since percentages were averages across several mice, a combination of variance in the placement of the slice across the preBötC would lead to different counts of subtypes if they were not distributed uniformly throughout the preBötC, and natural variability in the percentages of inhibitory neuronal subtypes across mice likely accounts for this effect. For semi-quantitative assessment of SST expression in preBötC and BötC regions a Histo score (H-score) was calculated from the data using the following formula based on manufacturer’s Data Analysis Guide (Advanced Cell Diagnostics):

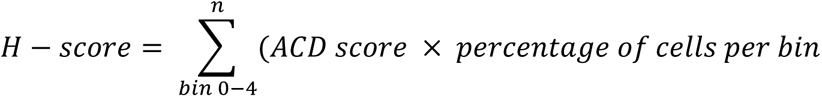

### Immunohistochemistry

After the completion of the experiment, mice were deeply anesthetized with an intraperitoneal injection of sodium pentobarbital (50mg/kg) and perfused through the ascending aorta with saline followed by 4% paraformaldehyde (PFA) in phosphate buffered saline (PBS). The brainstem was then dissected, postfixed overnight in 4% PFA at 4^°^C, and cryoprotected by equilibration in 30% sucrose in PBS at 4^°^C for 24-48 hours before sectioning. Coronal or sagittal sections (40 μm) of the brainstems were then cut using a cryostat (CryoStar™ NX70, Thermo Scientific) and stored at 4^°^C until further processing. Sections were then stained with the following antibodies: rabbit α-Neuronal Nuclei (NeuN) (EMD Millipore, ABN78), chicken α-Green Fluorescence Protein (GFP) (Aves Labs, Inc., GFP-1010), and goat α-Choline Acetyltransferase (ChAT) (EMD Millipore, AB144P) in accordance with the standard free-floating immunohistochemical protocol (Bachman, 2013).

### Studies in anesthetized mice

#### Breathing recording

Mice were anesthetized with isoflurane, placed on a heating pad, and cannula(s) were connected to optic fiber(s). Anesthetic depth was varied between 0.5 and 4% via a nose cone and breathing was monitored by using a pressure transducer (Model DP103-10-871, Validyne Engineering Corporation) connected to a carrier demodulator (Model CD15-A-2-A-1, Validyne Engineering Corporation). Airflow was used to monitor breathing and to estimate respiratory variables.

Data for PRCs and cophase plots were acquired by delivering 100 ms pulses into the preBötC at random phase of the breathing cycle and at ≥10 s intervals, e.g., in Figure 2A. Individual mice were tested on several different days to establish reliability of the response (n=7 mice).

#### Extracellular recording

Glass micropipettes were pulled from borosilicate glass (OD 1.5 mm, ID 1.12 mm) on a horizontal puller (Sutter Instrument Co., Model P-97). The tip of the micropipette was coated with a fluorescence dye (Vybrant® Dil Cell-labeling solution, Life Technologies, reference V22885, MW: 933.88), filled with artificial cerebrospinal fluid (ACSF; 124 mM NaCl, 3 mM KCl, 1.5 mM CaCl_2_, 1 mM MgSO_4_, 25 mM NaHCO_3_, 0.5 mM NaH_2_PO_4_, and 30mM D-glucose), and combined with an optic fiber (200 μm, Doric Lenses or ThorLabs), which was placed 500-1000 μm above the tip of the electrode. Transfected mice were placed in the stereotaxic frame as described above. A window was drilled through the skull of sufficient size to allow a free range of movement to the combined electrode/fiber. The recording electrode was connected to a headstage (Siskiyou) and the signal amplified (Grass Model P511, Grass Instruments) and sampled at 20 kHz (PowerLab 16SP, ADInstruments). Respiratory-modulated units were found 4.6 to 4.9 mm below the cerebellar surface and 1.1 to 1.3 mm lateral to the midline. Post-mortem, the placement of the combined electrode/fiber was confirmed before immunostaining to preserve the dye using the fluorescence microscope (Axioplan2 Imaging, Carl Zeiss).

### Studies in freely moving mice

#### Breathing Recording

Adult male mice implanted with fiberoptic cannula(s) that were connected to a 473/593 nm dual wavelength laser (OptoDuet Laser, IkeCool) via optical fibers and a rotary joint (Doric Lenses) were placed in a whole body mouse plethysmograph (PLY4211, Buxco Research Systems, Sharon, CT), where the animal could move freely, with access to water and chow. Plethysmograph was connected to a pressure transducer (Model DP103-10-871, Validyne Engineering Corporation) and a carrier demodulator. Whole Body Plethysmography (WBP) was used to monitor breathing and to estimate respiratory variables.

#### Normoxia, hypoxia, and hypercapnia

We exposed awake behaving mice to hypoxia and hypercapnia; both known to increase respiratory network drive while eliciting distinct ventilatory response. The ventilatory response to hypoxia has been shown to be biphasic (Fung *et al*., 1996; Teppema and Dahan, 2010) (Mortola, 1996), i.e., an initial adaptive increase in breathing (Phase 1, within 1-2 min) followed by respiratory depression (Phase 2, 3-5 minutes). On the other hand, hypercapnia has been described as a potent breathing stimulant (Guyenet and Bayliss, 2015; Del Negro, Funk and Feldman, 2018). Responses to hypoxia and hypercapnia were first characterized using a control group consisting of mice virally-injected and implanted with fiber-optic cannulas exposed to hypoxia (8% O_2_, 5 minutes) and hypercapnia (5% CO_2_, 5 minutes) without photostimulation (5 mice, Figure S1 and Table S2). These respiratory perturbations were then used to study how SST-mediated modulation is affected by two additional behavioral states of increased excitability but with distinct ventilatory responses, i.e., depressed *f* (hypoxia-induced, Phase 2) and stimulated *f* (hypercapnia-induced). In the test group, 473 nm light was delivered to freely moving mice under normoxic conditions (21% O_2_) and during hypoxia and hypercapnia. 1 s continuous pulses were delivered every 60 s (regardless of phase), during normoxic, hypoxic, and hypercapnia conditions. Phase onset times of stimuli were determined during post hoc analysis. Stimulation parameters, e.g., onset, duration, and frequency, were controlled by LabChart Stimulator.

### Data analysis and statistics

Plethysmograph and airflow traces were analyzed in LabChart 7 (ADInstruments). Raw data was first filtered (digital high-pass filter, cut-off frequency of 1 Hz) and smoothed using a Triangular (Bartlett) window. Inflection points were detected using a peak detection algorithm based on a general shine shape and a minimum peak height. Pulse onset was also identified using a peak detection algorithm, which was based on a predefined threshold. Data was imported into R (R: The Project for Statistical Computing, The R Foundation) where all the analyses and statistical tests were implemented using a custom R code. Statistical significance was set at p<0.05.

#### Cophase plots

We adapted the concept of phase resetting described by (Paydarfar, Eldridge and Kiley, 1986; Winfree, 1987) to our experimental paradigm. Phase of photostimulation (ϕ) was defined as the phase of the breathing cycle at which the pulse began, i.e., latency from inspiration onset to pulse onset, and cophase (θ) as the latency from pulse onset to the subsequent cycle (Figure 2B). Phase and cophase were normalized by the period of the preceding cycle and shown as cycle units, i.e., 1 is the period of an unperturbed cycle before photostimulation (ϕ+θ=1). Phase-dependent effects of photostimulation on the respiratory cycle were investigated via cophase plots, i.e., ϕ against θ. Deviations from the unity line (ϕ=-θ+1) indicate changes in respiratory cycle duration, such as prolongation (ϕ+θ>1), shortening (ϕ+θ<1), or resetting (θ≈0) of the cycle.

#### Cycle duration vs. baseline *f*

Normalized duration (ϕ+θ) of photostimulated and unperturbed, i.e., without photostimulation, breathing cycles was plotted against the baseline *f*. We used the duration of unperturbed cycles as a control to establish the range of 95% (mean±2SD) confidence. A photostimulated cycle was deemed to be significantly perturbed, i.e., shortened or prolonged, if its duration fell outside this confidence interval. Based on these significantly perturbed cycles, we identified the baseline *f* threshold at which the bimodal effect on cycle duration became very rare (<0.5%). The density of the normalized cycle duration and cophases were then investigated for *f*<2.5 Hz and *f*≥2.5 Hz.

#### Phase Response Curves

PRCs were investigated by plotting ϕ against each respiratory-related variable: inspiratory duration (T_I_), Amplitude (Amp), and expiratory duration (T_E_). For photostimulated and unperturbed breathing cycles, we computed the boxplots for T_I_, Amp, and T_E_ for each phase bin (0.1) with baseline *f*<2.5 Hz or ≥2.5 Hz. To determine statistical significance, we used unpaired t-tests between the respiratory cycle preceding photostimulation (Pre) and the photostimulated cycle (Stim). We calculated the percentage of breaths with a significant increase or decrease in T_E_ compared with the unperturbed breaths (mean±2SD), upon photostimulation at different phases of expiration (bin=0.1) and upon photostimulation at different baseline *f* (bin=0.2). To take into account differences in the baseline, each variable was also normalized by its counterpart during the preceding respiratory cycle, i.e., T_I_ normalized, Amp normalized, and T_E_ normalized and plotted in PRCs. Additionally, we calculated the percentage of unperturbed or photostimulated cycles per 0.2 bin of the normalized variable.

#### Hypoxia vs. Hypercapnia without photostimulation

Hypoxia stimulates breathing but when it is sustained, a decrease in the response is often observed (Vizek, Pickett and Weil, 1987; Voituron *et al*., 2009). The control group we tested showed (Figure S1 and Table S2), indeed, a significant increase in *f* during the first minute in hypoxia. After the second minute, however, *f* was significantly decreased. During hypercapnia, *f* was generally increased. To determine statistical significance of changes in *f* caused by hypoxia or hypercapnia, we used ANOVA and post-hoc Tukey multiple comparison tests. Further analysis in the test group were restricted to the last 3 minutes of each challenge when *f* was decreased in hypoxia and increased in hypercapnia.

#### Respiratory measurements in behaving mice

Baseline breathing frequencies in normoxia, hypoxia, and hypercapnia were calculated without photostimulation (Pre, 1 s). Breathing frequencies were then calculated during photostimulation (Stim, 1 s) in normoxia, hypoxia, and hypercapnia and compared to *f* without photostimulation. Moreover, breathing frequencies were calculated during the second following photostimulation (Post, 1 s) to verify if changes on breathing frequency were transient or lasted after termination of the pulse.

For each breathing cycle we calculated its a) Amp, b) latency, i.e., time between stimulus onset and the subsequent inspiratory peak, and c) duration, in normoxia, hypoxia, and hypercapnia. Cycles happening before photostimulation and occurring during photostimulation were selected, analyzed, and compared. Coefficients of variation of Amp, latency, and duration were calculated for Pre and Stim. To determine statistical significance among groups, we used ANOVA and post-hoc Tukey multiple comparisons.

### Image visualization and transfection quantification

Immunofluorescence staining was visualized and images were collected by confocal laser scanning microscopy (Zeiss LSM710). Multichannel (488, 570, and 647 nm) z-stacks were processed with ImageJ (Abràmoff, Magalhães and Ram, 2004; Schneider, Rasband and Eliceiri, 2012).

Using the nucleus ambiguus (NA), the facial nucleus (7N), and the Lateral Reticular nucleus (LRt) as anatomical landmarks, sagittal and coronal sections of SST-cre mice injected with AAV2/1-ChR2-eYFP were selected to include the core of preBötC (Figure 1A). Using a total of 11 sections (7 mice), either coronal (n=5 slices) or sagittal (n=6 slices), we estimated the transfection rate within a representative region of the preBötC (250 μm x 250 μm). Within this region, SST-ChR2-eYFP^+^ and ir-NeuN^+^ neurons were identified and quantified using the ImageJ plugin “Cell Counter”. In these sections, we estimated expression of eYFP^+^ in 15±6% of the preBötC neurons (SST-ChR2-eYFP^+^/ NeuN^+^). We confirmed also, and as previously characterized in (Tan *et al*., 2010), that preBötC SST-ChR2-eYFP^+^ neurons projected to the contralateral (non-injected) preBötC, and the ipsi- and contralateral Bötzinger Complex, BötC. Injections and optical cannula location were optimized to maximize transfection and photostimulation of preBötC and minimize surrounding areas such as the BötC.

**Figure 1.**
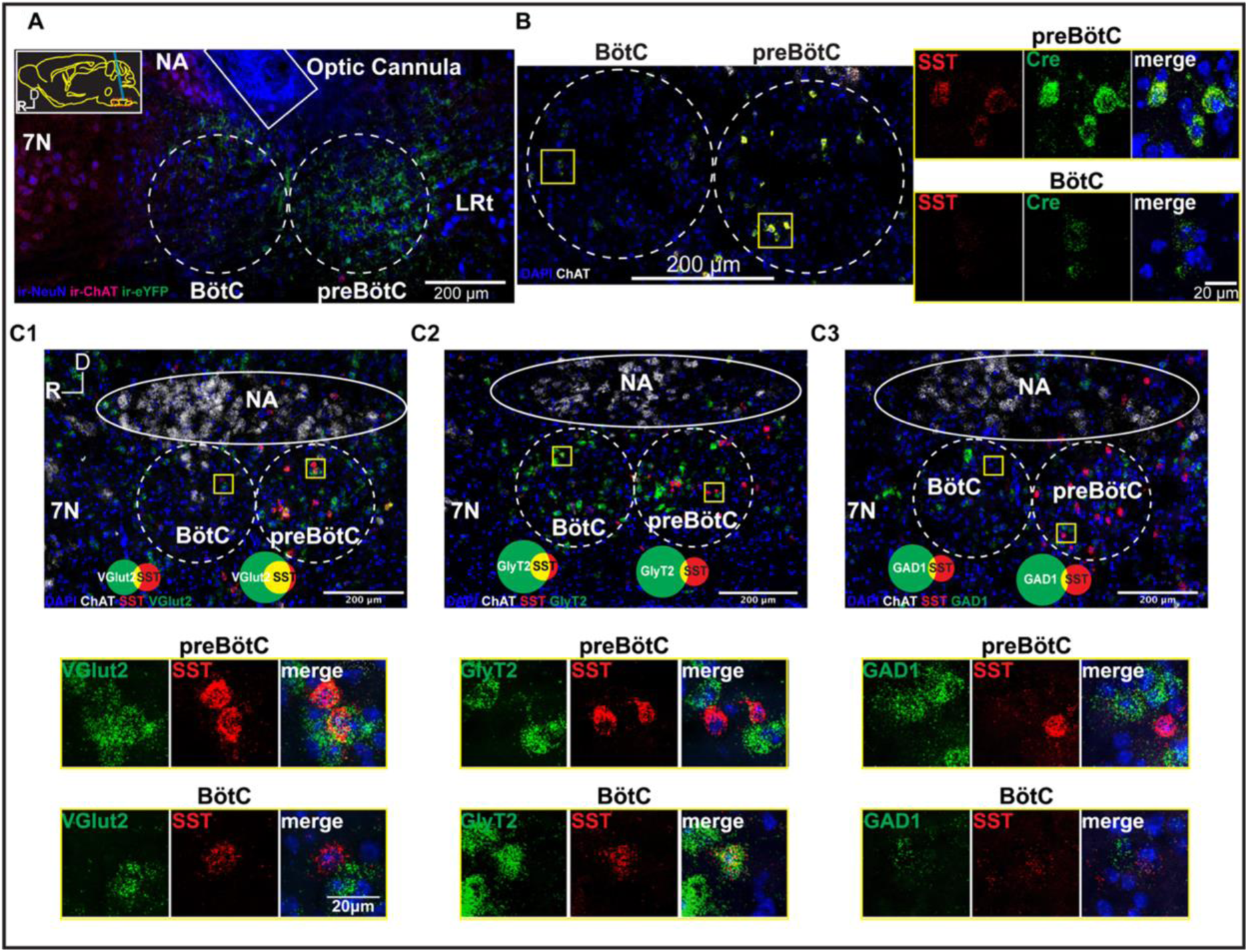
**A)** Sagittal section of medulla (see inset; D: dorsal; R: rostral) showing injection site and optical cannula track of a SST-cre mouse unilaterally injected with cre-dependent virus to selectively transfect preBötC SST^+^ neurons with the excitatory opsin ChR2 (ir-eYFP^+^, green). Ir-NeuN^+^ (blue) and ir-ChAT (magenta). **B) Left-** Sagittal section used for *in situ* hybridization RNA analysis (RNAscope). DAPI (blue) and ChAT (white). **Right-**mRNA expression of SST (red; left panels) and Cre (green; middle panels) in preBötC and BötC neurons. Merged channels shown in the rightmost panels. **C) Top**- Sagittal sections used for RNAscope analysis. DAPI (blue), ChAT (white), SST (red), VGluT2/GlyT2/Gad1 (green). **Bottom-** mRNA co-expression of SST (red; middle panels) and **C1)** VGluT2 (green; left panels), **C2)** GlyT2 (green; left panels), **C3)** GAD1 (green; left panels) in preBötC and BötC. Merged channels shown on rightmost panels. NA-Nucleus Ambiguus, 7N-Facial Nucleus, BötC- Bötzinger Complex, LRt- Lateral Reticular nucleus, preBötC- preBötzinger Complex.

## Results

### The majority of SST^+^ neurons are glutamatergic in preBötC but glycinergic in BötC

We virally transfected preBötC neurons to express ChR2 in transgenic mice with cre-expression driven by the SST promoter, i.e., SST-cre mice (Taniguchi *et al*., 2011). We confirmed that transfected neurons (15±6% of all preBötC neurons, n=11 slices, 7 mice) were mainly located in the preBötC immediately ventral to NA and that the optic cannula was properly positioned (Figure 1A). In order to interpret and understand the physiological role of these neurons, we first investigated the phenotypic identity of SST^+^ neurons in preBötC and in BötC (Figure 1B-C) of SST-cre mice. Using *in situ* hybridization RNA analysis (RNAscope), we confirmed that within preBötC and BötC all neurons that expressed SST RNA, i.e., SST^+^, co-expressed cre RNA, i.e., Cre^+^ (Figure 1B). Conversely, all Cre^+^ neurons expressed SST RNA; however, preBötC SST^+^ neurons had a significantly higher RNA SST expression than BötC SST^+^ neurons (H-score 282±4.2 vs. 227±20, n=3 mice each, p=0.03; see METHODS).

In preBötC, the SST^+^ subpopulation was 35% (60/173, here and in all subsequent RNAscope results, data are from three 20 μm sections from three mice) of the glutamatergic (vGluT2^+^) neuron population. Most preBötC SST^+^ neurons expressed the transcript for vGluT2 (vGluT2^+^SST^+^; 87% (60/69), Figure 1C1). When we separately examined co-localization of SST transcripts with inhibitory markers in preBötC, 21% (15/71) were glycinergic (GlyT2^+^SST^+^; Figure 1C2) and 15% (10/68) GABAergic (GAD1^+^SST^+^; Figure 1C3). Note that percentages exceeded 100%, which was due to variability between tissue samples (see METHODS) and that GAD1 and GlyT2 were co-expressed in some neurons (Koizumi *et al*., 2013; Rahman *et al*., 2013, 2015). In BötC, SST^+^ neurons were mostly glycinergic (SST^+^GlyT2^+^, 77% (50/65), Figure 1C2); 25% (13/51) were glutamatergic (SST^+^vGluT2^+^, Figure 1C1) and 25% (13/52) GABAergic (SST^+^GAD1^+^, Figure 1C3).

### State- and phase-dependent breath by breath modulation by preBötC SST+ neurons, in intact anesthetized mice

During normal rodent behavior, various parameters of breathing pattern, e.g., cycle duration, *f*, tidal volume, can vary rapidly and considerably, even within a breath, (Kabir *et al*., 2010), e.g., due to epochs of sitting still, grooming, chewing, walking. In order to reduce these confounds, we first investigated breath by breath modulation by preBötC SST^+^ neurons in anesthetized, but otherwise intact, mice across different phases of the breathing cycle. We identified changes in the cophase (θ), i.e., the interval from stimulus onset to the subsequent inspiratory onset normalized by the period of the preceding unperturbed breath, and other breathing-related variables, i.e., T_I_, expiratory duration (T_E_), and Amp, when 100 ms pulses excited ChR2 transfected preBötC SST^+^ neurons, e.g., Figure 2A (see METHODS).

**Figure 2.**
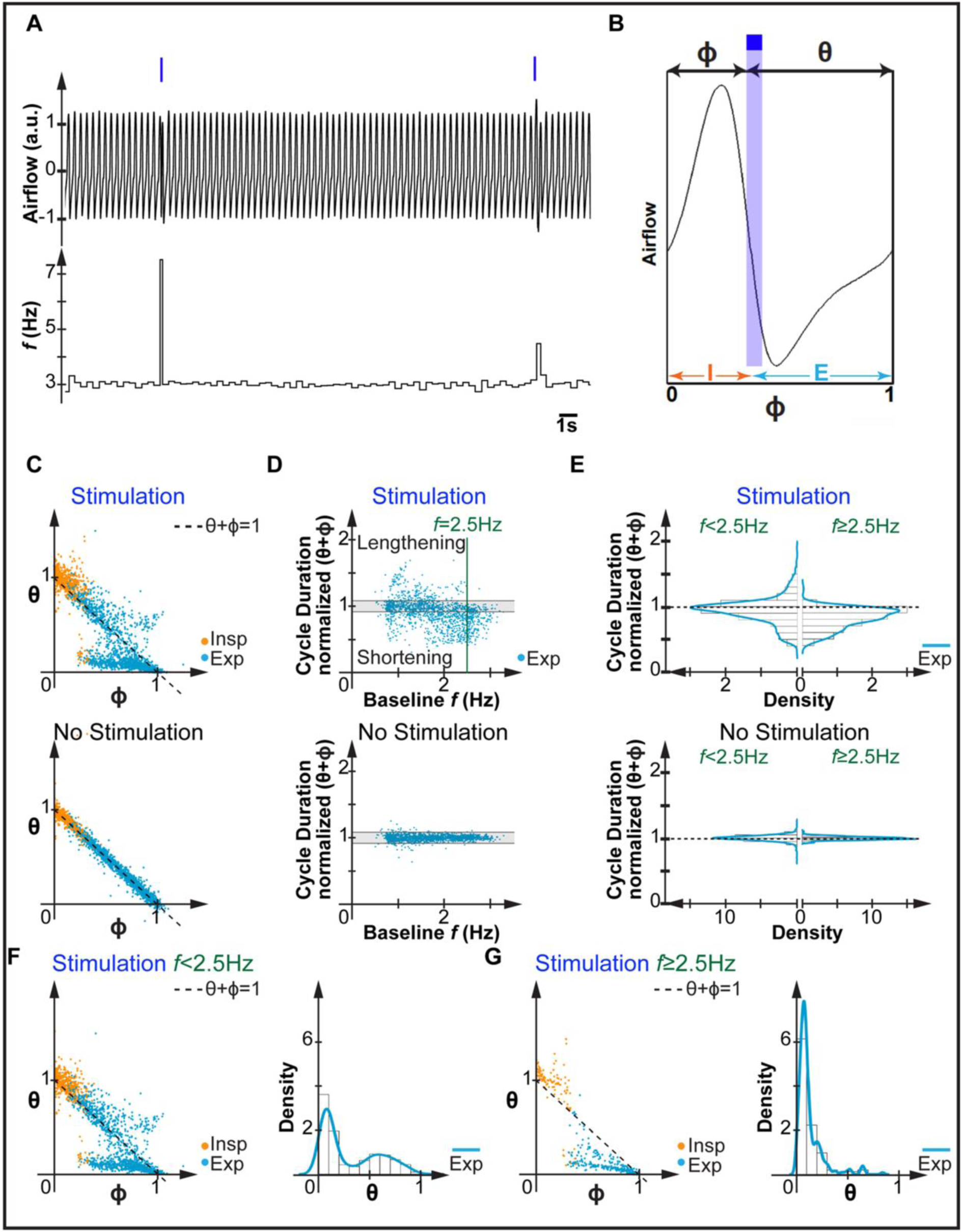
**A)** Representative traces of airflow (**Top**) and breathing frequency (*f*, Hz; **Bottom**) of an anesthetized mouse receiving bilateral photostimulation (473 nm light pulse) of preBötC SST^+^ neurons. **B)** Airflow trace for one respiratory cycle that includes inspiration (I, orange) and expiration (E, azure). For each pulse (blue box), phase of photostimulation (ϕ) and cophase (θ) were calculated and plotted in the cophase plots (ϕ vs. θ). **C) Top**- Phase of photostimulation (ϕ) vs. cophase (θ) when preBötC SST^+^ neurons were activated by 100 ms pulses delivered at different phases of inspiration (Insp, orange) or expiration (Exp, azure) in anesthetized mice (n=7). **Bottom-** Cophase plot (ϕ vs. θ) of unperturbed breaths, i.e., without photostimulation. Black dashed line indicates unity line if there is no effect or perturbation on duration of breathing cycle, i.e., ϕ+θ=1. **D) Top-** Normalized cycle duration (ϕ+θ) against baseline *f* when pulses were delivered during Exp. Green line indicates *f*=2.5 Hz and shaded area is range that includes 95% of the unperturbed breaths. **Bottom-** Normalized cycle duration (ϕ+θ) against baseline *f* for unperturbed breaths. Shaded area is the range that includes 95% of the unperturbed breaths. **E) Top-** Distribution of normalized cycle duration upon photo stimulation during Exp when *f*<2.5Hz (left panel) or ≥2.5Hz (right panel). **Bottom-** Distribution of normalized cycle duration of unperturbed breaths when *f*<2.5Hz (left panel) or ≥2.5Hz (right panel). **F-G)** Cophase plot in A in different ranges of *f*: *f*<2.5Hz (**F;** left panel) and *f*≥2.5Hz (**G;** left panel). Black dashed line indicates unity line if there is no effect or perturbation on duration of breathing cycle. Distribution of cophase (θ) upon photostimulation during expiration when *f*<2.5Hz (**F**; right panel) or *f*≥2.5Hz (**G**; right panel).

Generally, photostimulation of preBötC SST^+^ neurons during inspiration (Figure 2C top, orange points; 7 mice; compare with Figure 2C bottom) increased θ of individual cycles, i.e., prolonged breathing cycle duration (θ+ϕ>1). When stimulus onset was during expiration, however, the relationship between the phase of photostimulation and cophase was not linear. Instead, photostimulation during expiration either increased or decreased the cophase of individual cycles, i.e., prolonged (θ+ϕ>1) or shortened (θ+ϕ<1) cycle duration, respectively.

While we expected the response to photostimulation to have a phasic component dependent on the underlying rhythmic modulation of preBötC excitability, its bifurcation during expiration as a function of *f* was unexpected. We hypothesized that a difference in network excitability underlay the difference between these responses during expiration. To test this, we determined if responses parsed as a function of *f*, which is a surrogate measure of network excitability, i.e., higher the *f* the more excitable the network (Janczewski *et al*., 2013; Sherman *et al*., 2015; Baertsch, Baertsch and Ramirez, 2018). Consequently, we determined the effects of photostimulation during expiration on the cophase. We plotted the normalized cycle duration (θ+ϕ) against baseline *f* (Figure 2D top) vs. that in the preceding control, i.e., without photostimulation, cycle (Figure 2D bottom). We computed the 95% confidence interval (mean±2SD) of the control cycles (shaded area in Figure 2D bottom). A photostimulated breathing cycle was deemed significantly perturbed, i.e., shortened or prolonged, if its duration fell outside this confidence interval, which was the case in 59% of trial cycles vs. 5% (by definition) of control cycles. Of interest, these effects on cycle duration (θ+ϕ) were dependent on baseline *f*, with *f*=2.5 Hz the transition point between two distinct behavioral states (Figure 2E). When *f*≥2.5 Hz, photostimulation during expiration significantly shortened cycle duration in 70% of trials and significantly prolonged cycle duration in only 0.4% of trials. When *f*<2.5 Hz, photostimulation during expiration significantly shortened cycle duration in 42% of trials and significantly prolonged cycle duration in 16% of trials. When the cophase plot for all *f* (Figure 2C top) was revised for these distinct *f* ranges, the unimodal (*f*≥2.5 Hz, Figure 2G) and bimodal (*f*<2.5 Hz, Figure 2F) effects of photostimulation during expiration on the cophase became apparent.

Next, we investigated how photostimulation affected inspiratory (T_I_, Amp)- and expiratory (T_E_)-related variables when *f*<2.5 Hz and *f*≥2.5 Hz separately (Figure 3). Photostimulation during inspiration significantly increased the absolute (Figure 3A; Table S1) and normalized (Figure 4; see METHODS) values of T_I_ and Amp at any *f*, i.e., both *f*<2.5 Hz and *f*≥2.5 Hz. These effects were, however, more pronounced when *f*≥2.5 Hz, when photostimulation during inspiration also changed T_E_. Events with increased T_I_, Amp, and T_E_ were consistent with the observation of photostimulation-induced augmented/sigh-like events. When we investigated the effects of photostimulation during expiration on the absolute T_E_, we observed that photostimulation decreased the average T_E_ at any *f* (Figure 3B; Table S1). The effects were, however, more pronounced when *f*≥2.5 Hz and the pulse was delivered during early and middle expiration. Overall, the more excitable the network, i.e., higher *f*, the greater the impact of the photostimulation during inspiration and expiration.

**Figure 3.**
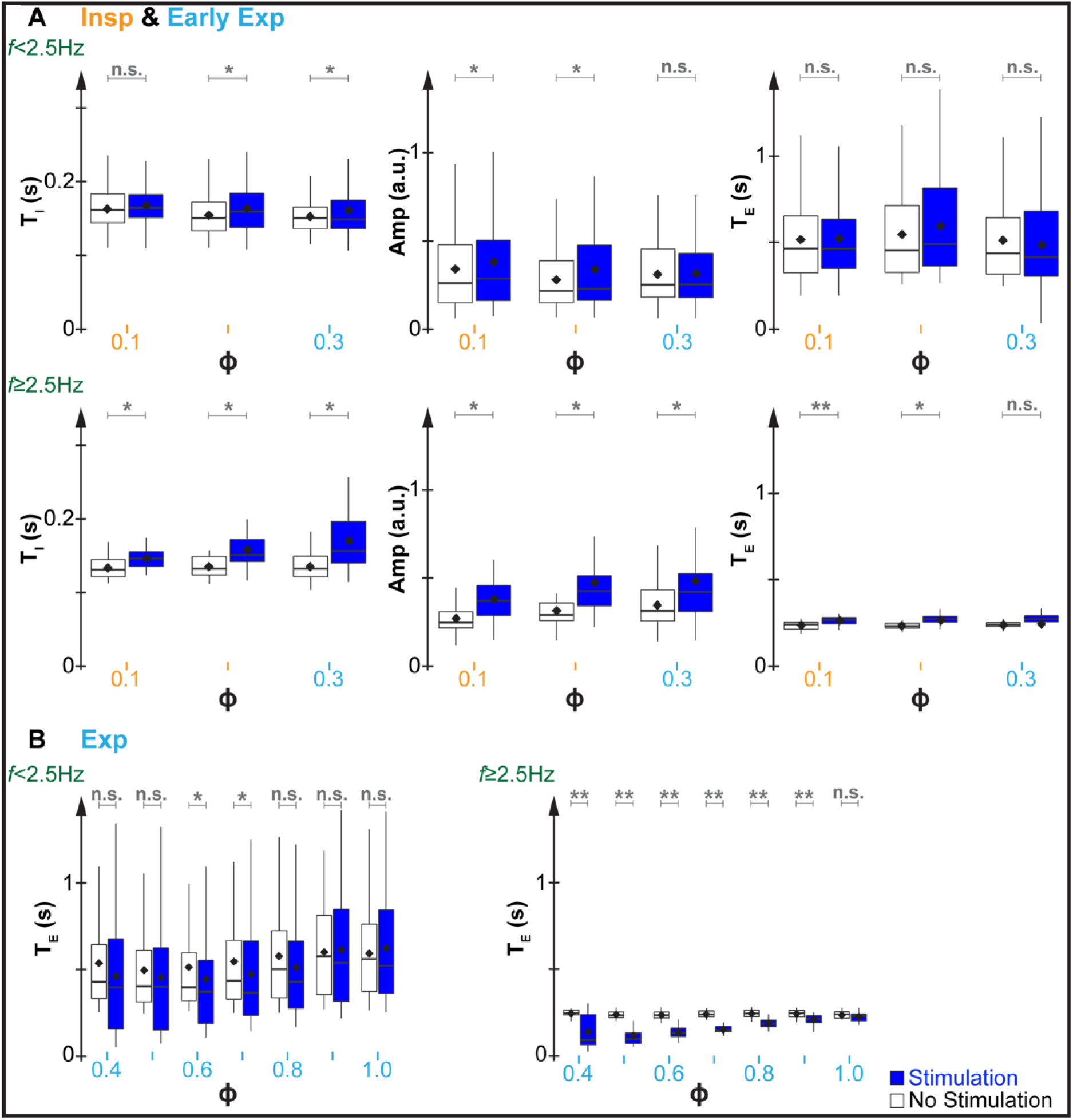
**A)** Photostimulation during inspiration (orange) and early expiration (azure): Phase Response Curves (PRCs) for T_I_, Amp, and T_E_ when *f*<2.5Hz (top panel) or *f*≥2.5Hz (bottom panel). **B)** Photostimulation during expiration: PRC for T_E_ when *f*<2.5Hz (left panel) or *f*≥2.5Hz (right panel). Black diamonds show means. Significance was found using unpaired t-tests. n.s. if p≥0.05, * if 10^−5^≤p<0.05 and ** if p<10^−5^.

**Figure 4.**
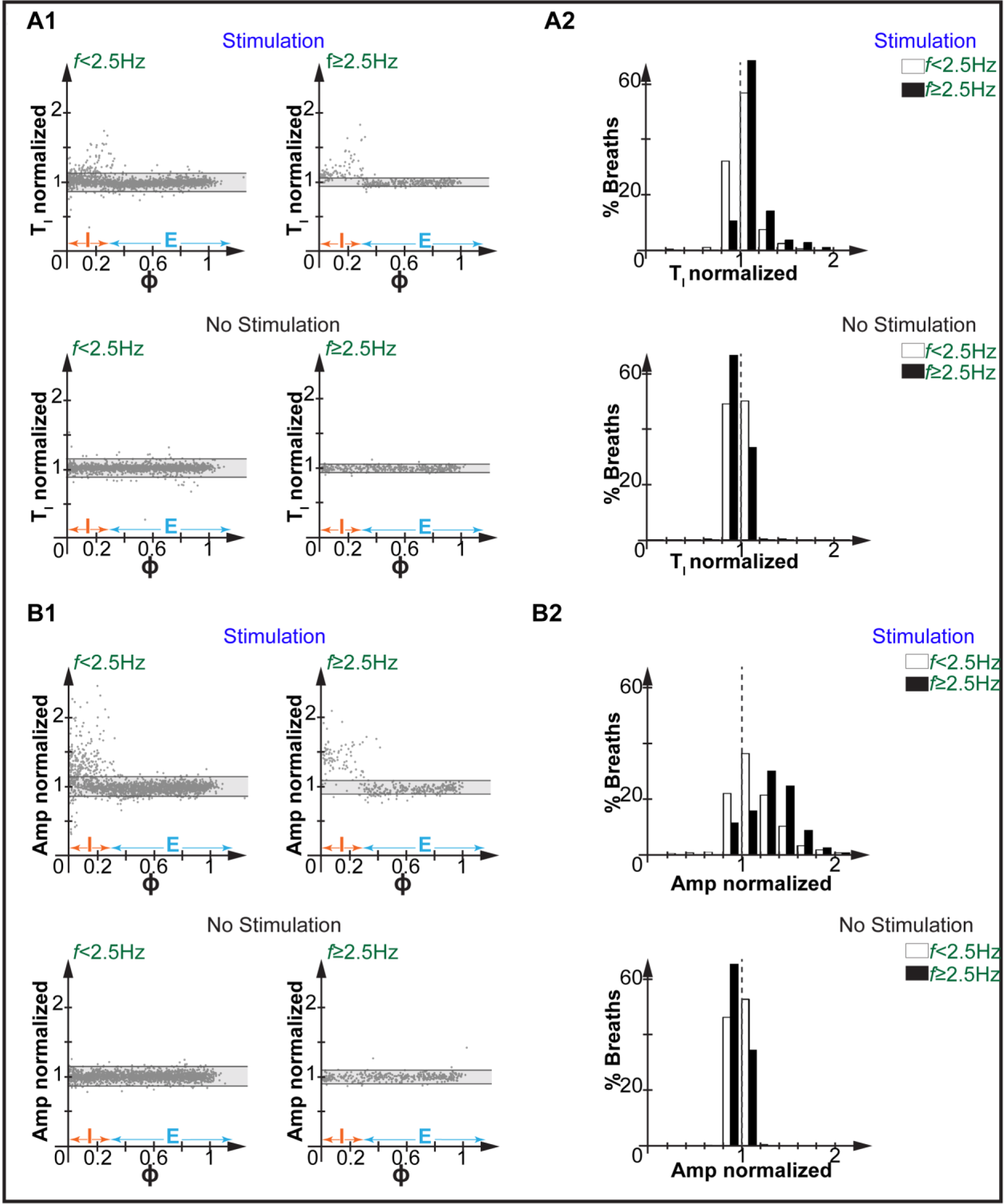
**A1**) PRC of normalized T_I_ with (top) or without (bottom) photostimulation when *f*<2.5Hz (Left) or *f*≥2.5Hz (Right); n=7. Shaded gray corresponds to the range within which 95% (mean±2SD) of unperturbed breaths that fell inside. I (orange) and E (azure) represent estimated inspiration and expiration in unperturbed breaths. **A2)** Distribution of breaths (%) with normalized T_I_ bellow or above unity (gray dashed line). White when *f*<2.5Hz and black when *f*≥2.5Hz. **B1-B2)** normalized Amp.

While analyses of the absolute value of T_E_ indicated that photostimulation during expiration decreased average T_E_ at any *f*, PCRs of normalized T_E_ (Figure 5) revealed that photostimulation during expiration prolonged (14% of trials) or shortened (41% of trials) T_E_ when *f*<2.5 Hz. In contrast, when *f*≥2.5 Hz prolongation of T_E_ was rare (0.4% of trials) and shortening of T_E_ was even more frequent (81% of trials). We note that photostimulation did not always significantly impact T_E_. In 15% and 45% of the trials photostimulation did not significantly change T_E_ when *f*≥2.5 Hz or *f*<2.5 Hz, respectively.

**Figure 5.**
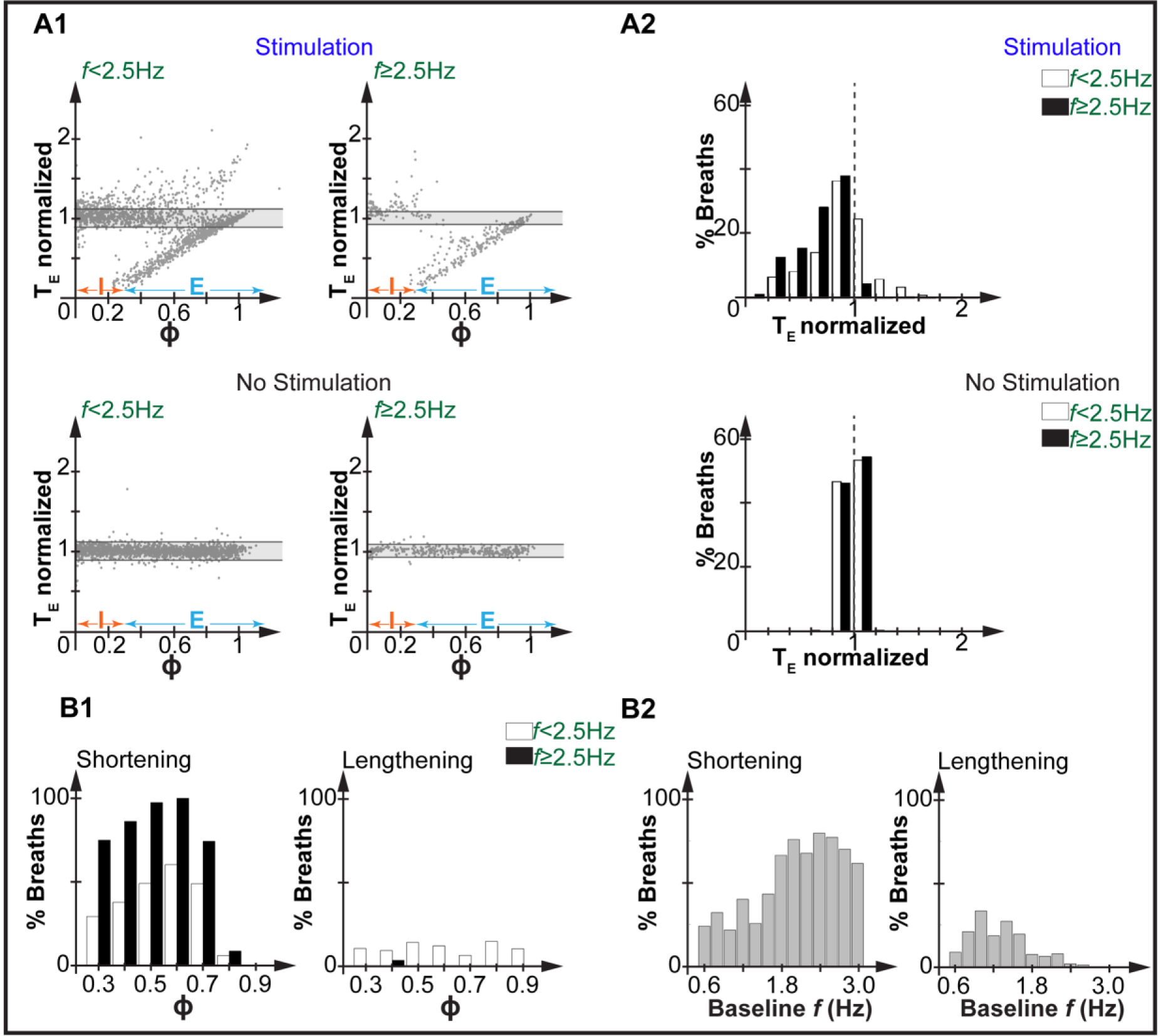
**A1**) PRC of normalized T_E_ with (top) or without (bottom) photostimulation when *f*<2.5Hz (Left) or *f*≥2.5Hz (Right); n=7. Shaded gray corresponds to the range within which 95% (mean±2SD) of the unperturbed breaths fell inside. I (orange) and E (azure) represent estimated inspiration and expiration in unperturbed breaths. **A2)** Distribution of breaths (%) with normalized T_E_ bellow or above unity (gray dashed line). White when *f*<2.5Hz and black when *f*≥2.5Hz. **B1)** Percentage of breaths with a significant decrease or increase in T_E_, i.e., shortening (left) or lengthening (right), respectively, upon photostimulation at different phases of expiration (bin=0.1). **B2)** Percentage of breaths with a significant decrease or increase in T_E_, i.e., shortening (left) or lengthening (right), respectively, upon photostimulation at different baseline *f* (bin=0.2).

To further characterize the relationship between phase of photostimulation and evoked response we normalized breathing variables (normalized T_I_, Amp, and T_E_) and plotted against the phase of photostimulation separately when *f*<2.5 Hz or *f*≥2.5 Hz.

Photostimulation during inspiration increased the incidence of breaths with higher normalized T_I_ and Amp in relation to the unity line (red vertical dashed line, Figures 4A2 and B2) at any *f*, with the effect being more pronounced when *f*≥2.5 Hz. Photostimulation during expiration, generally, increased the incidence of breaths with lower normalized T_E_ in relation to the unity line (red vertical dashed line, Figure 5A2), i.e., shortened T_E_, with the effect being slightly more pronounced when f≥2.5 Hz.

According to our postulated burstlet theory of breathing rhythm generation (Feldman and Kam, 2015; Ashhad and Feldman, 2020; Kallurkar *et al*., 2020), pre-inspiratory activity slowly increases as the respiratory cycle progresses, leading to initiation of a burstlet and ultimately a burst. Due to the considerable variability in the underlying network evolution with each breath (Carroll and Ramirez, 2013; Ashhad and Feldman, 2020), whether photostimulation at a particular onset time induces a phase advance or delay could vary from cycle to cycle. To assess that we plotted incidence of stimulation inducing phase advance (Figure 5B1, left plot) or phase delay (Figure 5B1, right plot) against the phase of photostimulation. Regardless of *f*, the probability of phase advance during expiration increased gradually until middle expiration (ϕ≅0.6-0.7), then rapidly declined to be very small or insignificant (Figure 5B1, left plot). Overall, the probability of phase advance was higher at all phases when *f*≥2.5 Hz. Similarly, the probability of phase advance increased as *f* increased (Figure 5B2, left plot). While the shortening effect on T_E_ occurred at any *f*, lengthening of T_E_ was only detected at lower *f* (Figure 5B2, right plot) and was independent of phase (Figure 5B1, right plot).

Finally, we monitored neuronal activity within the preBötC and confirmed the increased preBötC neuronal activity when photostimulation during inspiration triggered events with increased T_I_ and Amp (Figure 6A1) or with just increased T_I_ (Figure 6A2); both often happening in the same animal. Moreover, in the same mouse, photostimulation during expiration could have minor effects (Figure 6B1), delay (Figure 6B2) or advance (Figure 6B3) the phase. In all cases, we observed an increase in preBötC neuronal activity upon photostimulation.

**Figure 6.**
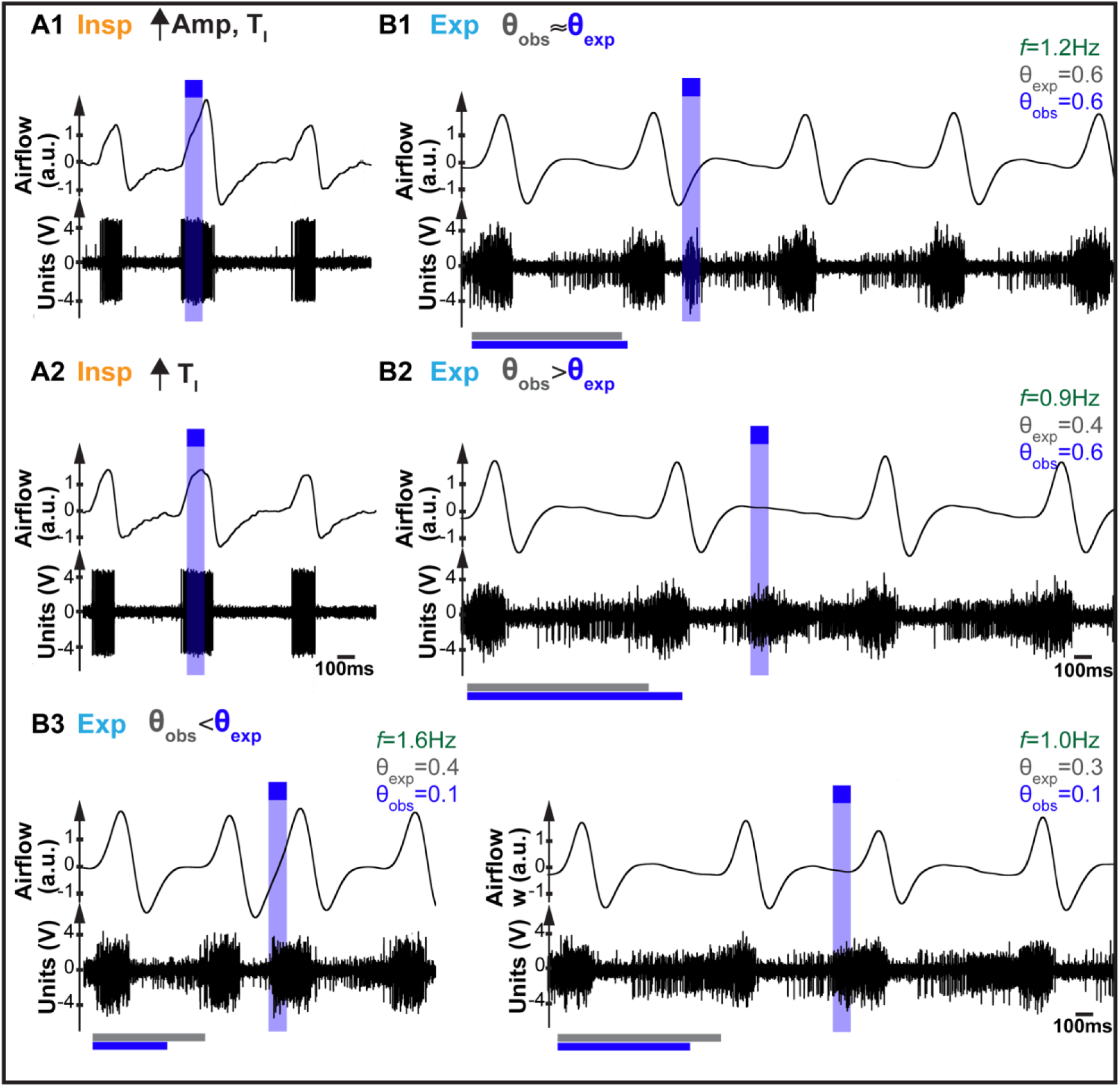
**A)** preBötC inspiratory single unit activity and motor output (airflow) when pulses (blue box) were delivered during inspiration to the same mouse. **A1)** Photostimulated breath with increased Amp and T_I_. **A2)** Photostimulated breath with increased T_I_, but not Amp. **B)** preBötC population activity and motor output (airflow) when pulses were delivered during expiration to the same mouse. **B1)** Examples of photostimulated breaths with little changes on the phase, i.e., θ_obs_ ≈ θ_exp_, **B2)** phase delay, i.e., θ_obs_ > θ_exp_, and **B3)** phase advance, i.e., θ_obs_ < θ_exp_. Phase of photostimulation (ϕ), observed cophase (θ_obs_), expected cophase (θ_exp_), and baseline breathing frequency (*f*). Horizontal gray and blue lines represent period of the preceding breath and photostimulated breath, respectively.

### Similar rapid and slow modulation of breathing by preBötC SST+ neurons in freely moving mice exposed to room air, hypoxia, and hypercapnia

To better understand the role that preBötC SST^+^ neurons could play in modulating breathing during normal behavior, where resting, i.e., baseline, *f* and its variability are considerably higher (see below), we extended this study to unanesthetized, awake freely moving mice where control resting *f*>4 Hz. We observed that the responses to photostimulation were similar to those observed in anesthetized mice when *f*≥2.5 Hz. Brief (100 ms) photostimulation during inspiration could trigger an augmented breath/sigh-like event, e.g., Figure 7B, #, whereas photostimulation during expiration could advance the phase of the breathing cycle, e.g., Figure 7B, *.

**Figure 7.**
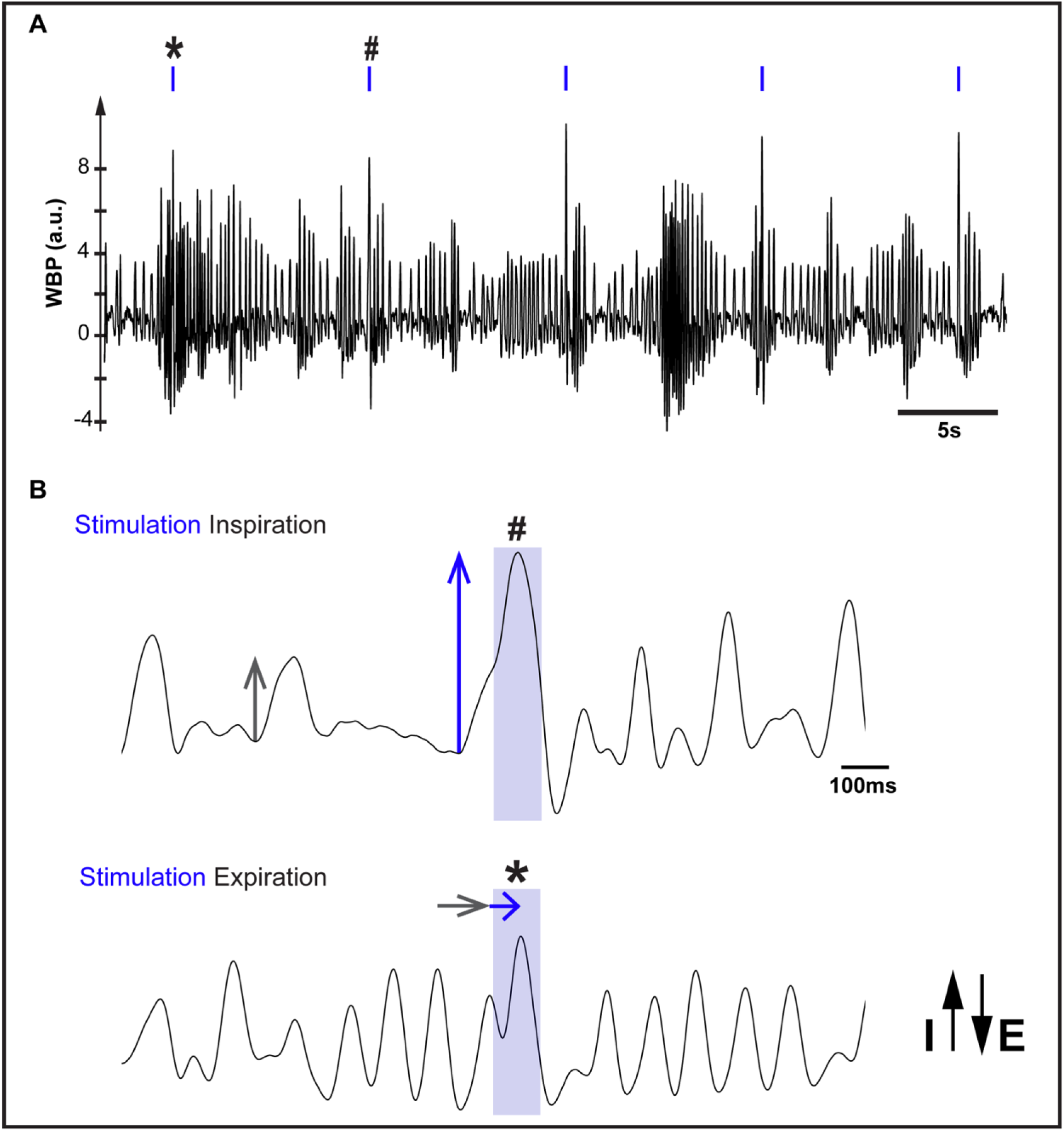
**A)** Representative whole body plethysmography (WBP) trace when 100 ms pulses (blue) were bilaterally delivered to a freely behaving mouse. Pulses * and # are shown in **B)** as examples of a pulse delivered during inspiration (top, #) and expiration (bottom, *). In WBP trace, upward represents inspiration (I) and downward expiration (E). Gray arrows indicate Amp (Top) and duration (Bottom) of the unperturbed breath prior to photostimulation. Blue arrows indicate Amp (Top) and duration (Bottom) of photostimulated breath.

Because of the considerable breath to breath variance in freely moving mice, normalized phase-based analysis (using short pulses) was not feasible. Instead, we looked at effects of longer pulses (1 s) on *f* and its variability, i.e., coefficient of variation (CV). Since 1 s photostimulation spanned several breathing cycles (up to 10, 4, and 9 breaths in normoxia, hypoxia, and hypercapnia, respectively), we investigated responses of the first four full breathing cycles after 1 s pulse onset, which we called photostimulated breathing cycles, and compared with 1 s of unperturbed breathing cycles immediately preceding photostimulation as control (see METHODS). Responses were then separated into two time domains (see (Cui *et al*., 2016)): i) within the first 100 ms of pulse onset, when the first photostimulated breathing cycle occurred, and ii) during the remaining duration of the pulse, i.e., 100 ms to 1 s, when the 3 subsequent photostimulated cycles we analyzed occurred (Figure 8A). For each photostimulated breathing cycle, we measured: i) time between stimulus onset and inspiratory peak, i.e., peak latency; ii) cycle duration, and; iii) peak inspiratory amplitude (Amp), and further explored the effects of hypoxia and hypercapnia (see METHODS) on these responses.

**Figure 8.**
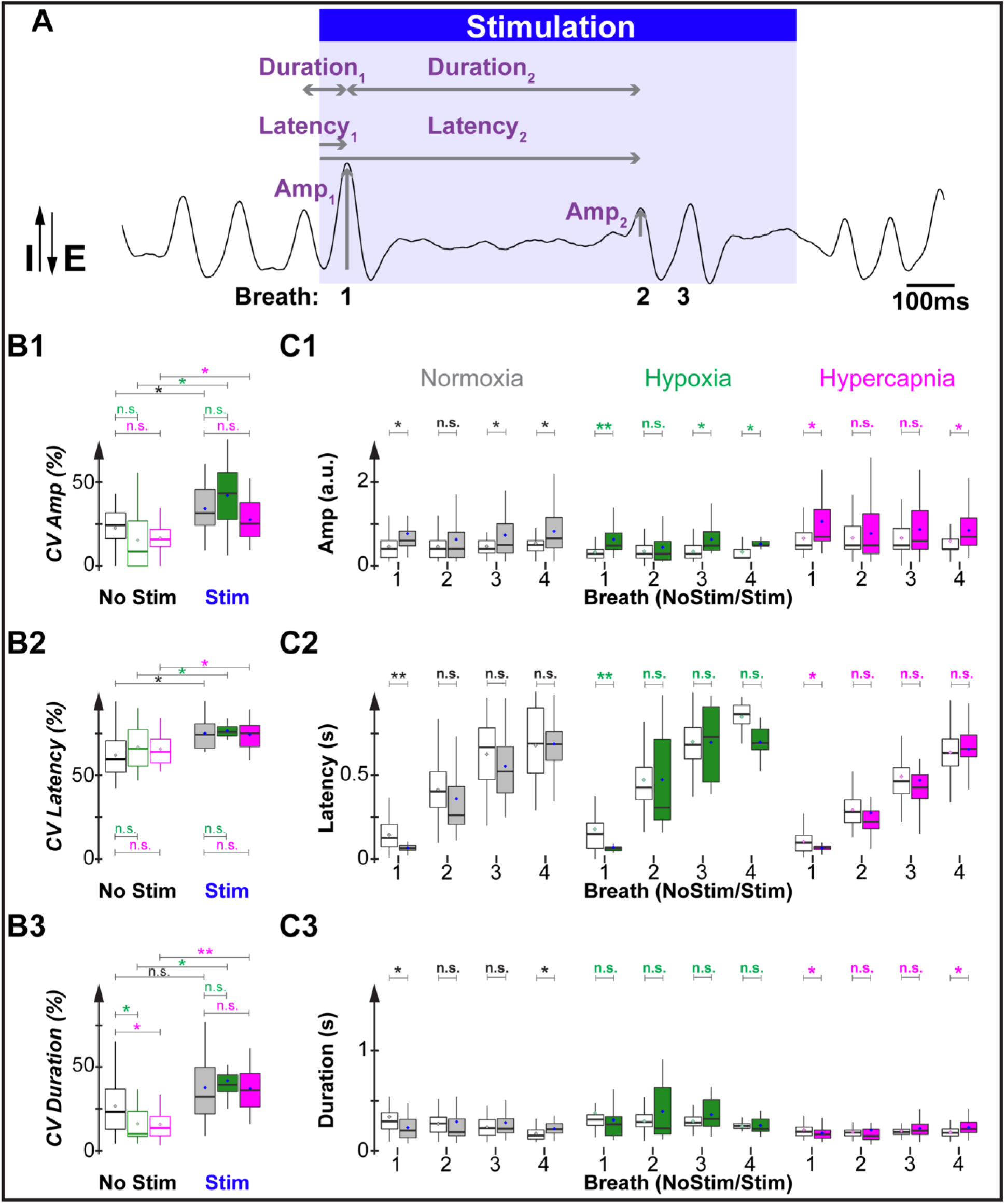
**A)** Sample WBP trace from a representative freely behaving mouse in normoxia when a 1s pulse (blue) was delivered. Vertical arrows indicate amplitude for the first (Amp_1_) and second (Amp_2_) photostimulated breaths following pulse onset. Horizontal single arrows indicate peak latency, i.e., delay time between stimulus onset and inspiratory peak, for the first (Latency_1_) and second (Latency_2_) breaths following pulse onset. Horizontal double arrows indicate duration of the first (Duration_1_) and second (Duration_2_) photostimulated breaths, i.e., time between inspiratory peaks. **B1, B2, and B3)** Coefficient of variation (CV) of breath Amp, peak latency, and duration with and without photostimulation, respectively. **C1)** Amp, C2) latency, and C3) duration for the first four breaths following pulse onset (Stim) and four unperturbed breaths preceding photostimulation (No Stim, white boxplots) in normoxia (left panel, Stim in gray), hypoxia (middle panel, Stim in green), and hypercapnia (right panel, Stim in magenta). Diamonds show means. Significance between unperturbed and photostimulated breaths was found using unpaired t-tests. Significance among respiratory conditions were found using ANOVA and post hoc Tukey’s multiple comparisons. n.s. if p≥0.05, * if 10^−5^≤p<0.05 and ** if p<10^−5^.

Both hypoxia and hypercapnia increase ventilation, an indication of an increase in overall respiratory network excitability, but hypoxia decreased *f* and hypercapnia increased *f* (see METHODS, Figure S1 and Table S2).

In absence of photostimulation, hypoxia and hypercapnia significantly changed *f* (normoxia: 4.4 ± 0.3 Hz; hypoxia: 3.4 ± 0.1 Hz; hypercapnia: 5.4 ± 0.2 Hz, Figure 9 and Table S3). During 1 s photostimulation, *f* decreased significantly in all conditions (normoxia + 1 s pulse: 3.3 ± 0.2 Hz; hypoxia + 1 s pulse: 2.0 ± 0.1 Hz; hypercapnia + 1 s pulse: 3.8 ± 0.3 Hz, Figure 9 and Table S3). As expected, in freely moving mice breathing variability was much higher compared to anesthetized mice. In normoxia, control, i.e., without 1 s photostimulation, breathing cycles had coefficients of variation of CV_Amp_: 23 ± 2%, CV_duration_: 29 ± 4%, CV_latency_: 62 ± 2%; Figures 8B1-D1 and Table S4). Then, still without photostimulation, the variability in duration was significantly reduced by hypoxia or hypercapnia (hypoxia: CV_duration_: 16 ± 2%; hypercapnia: CV_duration_: 16 ± 2%). In general, 1 s photostimulation of preBötC SST^+^ neurons increased breathing variability (Amp, latency, and duration) in normoxia (CV_Amp_: 34 ± 4%, CV_duration_: 41 ± 6%, CV_latency_: 77 ± 3%), hypoxia (CV_Amp_: 42 ± 8%, CV_duration_: 42 ± 5%, CV_latency_: 82 ± 4%), and hypercapnia (CV_Amp_: 28 ± 3%, CV_duration_: 44 ± 4%, CV_latency_: 74 ± 2%). In sum, effects of 1 s photostimulation on *f* and breathing variability with decreased (hypoxia-induced) or increased (hypercapnia-induced) *f* did not differ from normoxia.

**Figure 9.**
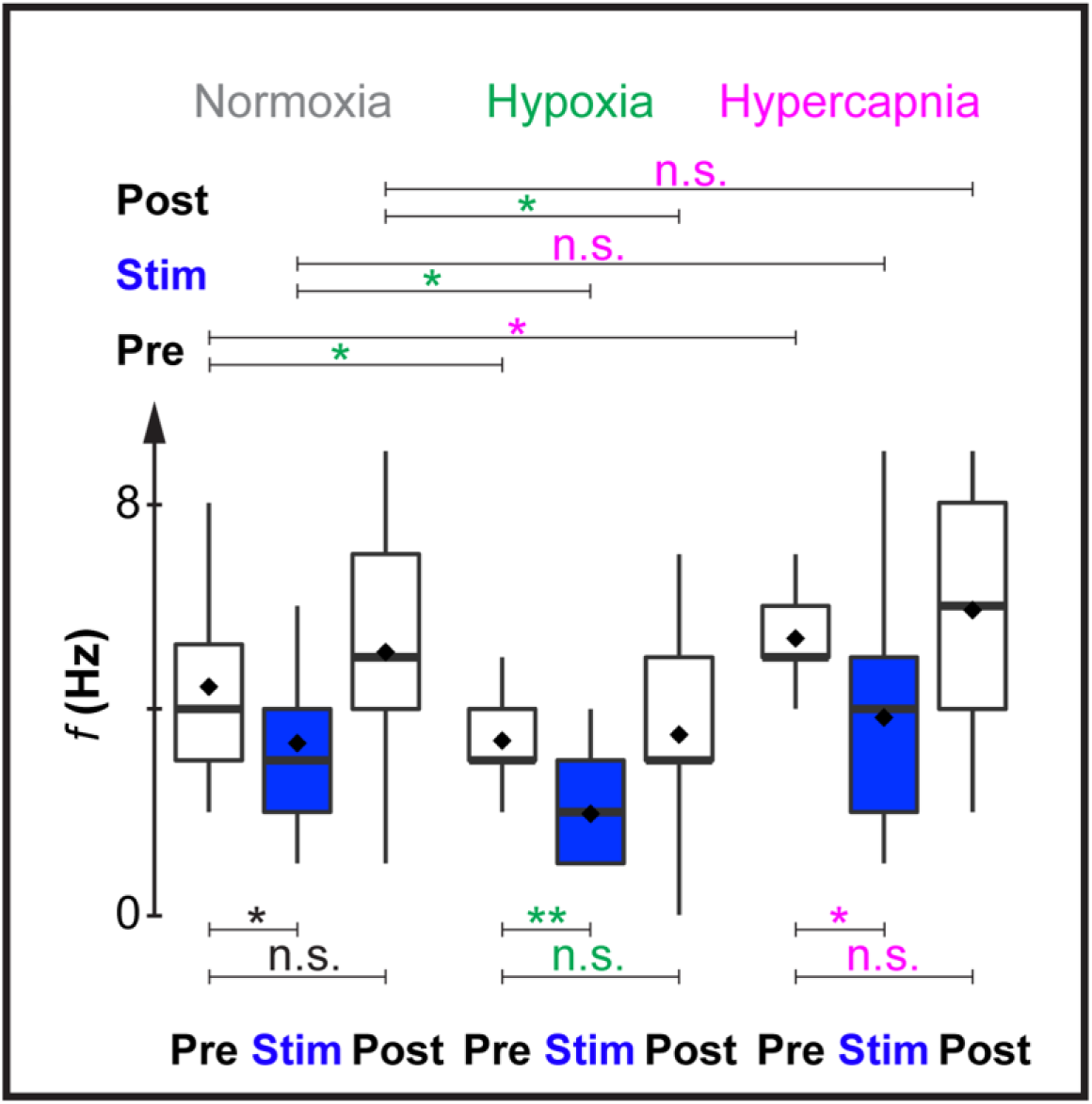
Breathing frequency *f* (Hz) prior (Pre, white), during (Stim) and after (Post, white) photostimulation (1 s bins) in normoxia (gray), hypoxia (green), and hypercapnia (magenta). Black diamonds represent mean. Significance among respiratory conditions or Pre, Stim, and Post bins were found using ANOVA and post hoc Tukey’s multiple comparisons. ns if p≥0.05, * if10^−5^≤p<0.05 and ** ifp<10^−5^.

We then compared responses of consecutive photostimulated breathing cycles with the unperturbed preceding control breathing cycles, irrespective of the phase of pulse onset (Figures 8C1, 8C2, 8C3 and Table S5). In normoxia, 1 s photostimulation significantly increased Amp of the first photostimulated cycle (Figure 8C1, Amp_1_) and decreased its latency (Figure 8C2, Latency_1_) and duration (Figure 8C3, Duration_1_). The inspiratory peak of the first photostimulated cycle occurred 69 ± 3 ms after pulse onset compared with 145 ± 13 ms in preceding control cycles (see METHODS); indicating that the first photostimulated cycle was shorter (230 ± 16 ms) than control cycles (337 ± 33 ms). Similar modulation of Amp and peak latency occurred in hypoxia (Figures 8C1 and 8C2), where the inspiratory peak of the first photostimulated cycle occurred 72 ± 6 ms after pulse onset in contrast to 179 ± 21 ms in unperturbed cycles. During hypercapnia, Amp (Figure 8C1) and peak latency (Figure 8C2) were similarly modulated. The inspiratory peak of the first cycle after pulse onset, i.e., first photostimulated cycle, had increased Amp that occurred after 67 ± 3 ms in contrast to unperturbed cycles after 104 ± 9 ms. In hypercapnia perturbed cycles were shorter (178 ± 8 ms) than unperturbed cycles (210 ± 10 ms). In brief, SST-modulation during the first hundred of milliseconds of the photostimulation in a depressed (hypoxia-induced) or stimulated (hypercapnia-induced) baseline *f* was identical to that observed in normoxia, i.e., increased Amp and decreased latency and duration.

During >100 ms to 1 s of the pulse, the majority of the breathing cycles had increased Amp in normoxia, hypoxia, and hypercapnia (Figure 8C1). One exception was the second cycle, which Amp dropped significantly. In all conditions, we did not observe a phase advance, i.e., decrease peak latency, in the subsequent cycles (Figure 8C2), but it prolonged the duration of the fourth cycle in normoxia and hypercapnia (Figure 8C3).

In summary, in different behavioral states, i.e., isoflurane-anesthetized mice with *f* < or ≥2.5 Hz, and awake, freely behaving mice in normoxia, hypoxia, and hypercania, preBötC SST^+^ neurons can modulate breathing in a phase- and state-dependent manner.

## Discussion

The mechanisms underlying generation of normal resting breathing patterns in mammals are complex and are highly state-dependent, in experimental conditions and in normal behavior. We investigated modulation of breathing pattern in several states in adult mice: i) anesthetized, but otherwise intact, and; ii) freely moving, exposed to room air, hypoxic or hypercapnic gas mixtures. The effects of activation of preBötC SST^+^ neurons on pattern generation were both phase- and state-dependent, with differences across states most likely the result of changes in bCPG network excitability.

### Phenotype of preBötC SST^+^ neurons

preBötC SST^+^ expressing neurons are largely glutamatergic (87% glutamatergic, 21% glycinergic, and 15% GABAergic; Figure 1C). [Note: This is a similar percentage of glutamatergic but a much higher percentage of inhibitory (glycinergic and/or GABAergic) neurons compared to a previous study (89% glutamatergic, 1% glycinergic; (Stornetta *et al*., 2003). This disparity could be due to differences between mouse and rat as well as differences in technique, as we used RNAscope with improved probes, signal amplification, and background noise suppression that have higher levels of sensitivity and selectivity compared to previous methods. Moreover, mRNA expression does not always reflect protein expression (de Sousa Abreu *et al*., 2009; Maier, Güell and Serrano, 2009)]. In contrast to the preBötC, in the immediately rostral BötC the majority of SST^+^ neurons were glycinergic (25% glutamatergic, 77% glycinergic, and 25% GABAergic). We recognize the possibility that some inhibitory preBötC or BötC SST^+^ neurons were also transfected and photostimulated, q.v., (Sherman *et al*., 2015; Baertsch, Baertsch and Ramirez, 2018), when we targeted glutamatergic preBötC SST^+^ neurons. The likelihood that BötC SST^+^ neurons were stimulated was minimized due to the small volume of the injections (<50 nL) within the preBötC and placement of the optical fiber over the preBötC, caudal to the BötC. We discuss the results of modulation of breathing by photostimulation of SST^+^ neurons with the explicit presumption that the dominant effects were due to activation of glutamatergic preBötC SST^+^ neurons.

### Rapid (≤100 ms) modulation

In isoflurane-anesthetized mice with *f* between 0.5-3.3 Hz, photostimulation of ChR2 transfected preBötC SST^+^ neurons at different phases of the breathing cycle produced a repertoire of changes in breathing pattern (Figures 2-6). With *f*≥2.5 Hz, photostimulation during: i) inspiration (ϕ=0-0.2) evoked augmented, i.e., sigh-like, breaths with increased T_I_, Amp, and T_E_ of the photostimulated cycle, and; ii) early-to-middle expiration (ϕ=0.3-0.8) advanced the respiratory phase, i.e., the next inspiration started sooner than expected. With *f*<2.5 Hz, the effects of photostimulation during inspiration were similar to those with higher *f*. However, during expiration there was a bimodal effect on T_E_, i.e., either an advance or delay in the onset of the next inspiration (Figures 2F and 5B1-B2). Phase advance occurred at all *f* and with a gradual increase in probability from early to middle expiration, when it reached its maximum (Figure 5B1, left plot). Photostimulation at ϕ=0.6-0.7 significantly shortened 100% of the trials when *f*≥2.5 Hz. The percentage of breaths with phase advance was notably less frequent when *f*<2.5 Hz. Photostimulation at the end of expiration (ϕ≥0.9) did not advance the phase, at any *f* (Figure 5B1, left plot). These results clearly reflect dynamic processes underlying expiration and are consistent with the fact that expiration is divided into i) an interburst interval (IBI), a quiescent phase of inspiratory motor activity following the termination of the inspiratory burst when the cycle is, presumptively, refractory to initiating another inspiratory burst, and; ii) preinspiration, which normally immediately precedes, and is hypothesized (Kam *et al*., 2013; Feldman and Kam, 2015; Ashhad and Feldman, 2020; Kallurkar *et al*., 2020) to initiate, each inspiratory burst. We propose that photostimulation of preBötC during expiration can “accelerate” the percolation of activity within the preBötC (Ashhad and Feldman, 2020) leading to premature initiation of the inspiratory burst. The contribution of preBötC SST^+^ neurons to the phase advance appears more pronounced during middle expiration, when the preinspiratory phase is, presumably, already in process; the effect is also more pronounced at higher *f*, either because: the expiratory phase is shorter, so that the 100 ms fixed duration pulse is more likely to overlap with preinspiratory activity, or; at higher *f*, excitability is higher with consequential greater effect on preinspiratory activity. Interestingly, only at lower *f*, photostimulation of preBötC SST^+^ neurons during expiration could also delay the phase, regardless of the phase of photostimulation (Figures 5B1, right plot; and 5B2, right plot). Importantly, we observed trials in the same mouse where photostimulation could induce phase advance (Figure 6B3) or phase delay (Figure 6B2) in neighboring trials, indicating that this effect was truly bimodal, and not due to intersubject variability, e.g., differences in transfection amongst mice.

What mechanism underlies these state-dependent responses during expiration? We suggest a possible schema in the context of our proposed “burstlet theory” (Kam *et al*., 2013; Feldman and Kam, 2015; Ashhad and Feldman, 2020; Kallurkar *et al*., 2020) (Figure 10). Under normal conditions, we propose that rhythmogenic burstlets trigger inspiratory bursts in each cycle (*ibid.*), effectively determining T_E_ (Figure 10A3 and B1). These burstlets are preceded by a period of percolating activity where random firing of rhythmogenic neurons becomes increasingly synchronized (Ashhad and Feldman, 2020). If we assume that there is a threshold for percolation to precipitate a burstlet (τ_burstlet_), and a second threshold for a burstlet to trigger a burst (τ_burst_), and that these thresholds are noisy but always exceeded during normal breathing, then photostimulation can lengthen or shorten T_E_ depending on whether activation can trigger percolation and/or burstlets to prematurely exceed their thresholds. Additionally, even though we controlled the magnitude of optogenetic stimulation, the effect of each stimulus would be modulated by the degree of ongoing synchrony (Ashhad and Feldman, 2020). First, we consider that the same optogenetic stimulus may or may not trigger a burstlet depending on the instantaneous threshold. If it does not reach threshold for burstlet generation, there should be no significant phase advance or delay, which we observed mostly during low *f* (Figure 10B2). When, however, photostimulation does trigger a burstlet, the burstlet may or may not reach its threshold for triggering a burst. If the burstlet does not reach threshold for burst generation, the percolating activity will be reset resulting in a consistent phase delay (Figure 10B3), and indeed, a consistent phase delay throughout the expiratory phase (Figure 5B1, right plot), but only under low *f* conditions (Figure 5B2, right plot). Since low *f* is associated with longer bursts, hence longer refractory periods (Janczewski *et al*., 2013), we also consider effects on the refractory period as a nonexclusive mechanism underlying phase delay. Distinctly to trials where photostimulation delayed the phase, a phase advance occurs if photostimulation evokes sufficient burstlet activity to trigger a burst. Phase advance was observed at all *f*, but was much more frequent when T_E_ was shorter, i.e., at high *f* (Figure 10A3) compared to low *f* (Figure 10B1). At high *f*, photostimulation during middle expiration always resulted in a phase advance (Figure 5B1, left plot).

**Figure 10.**
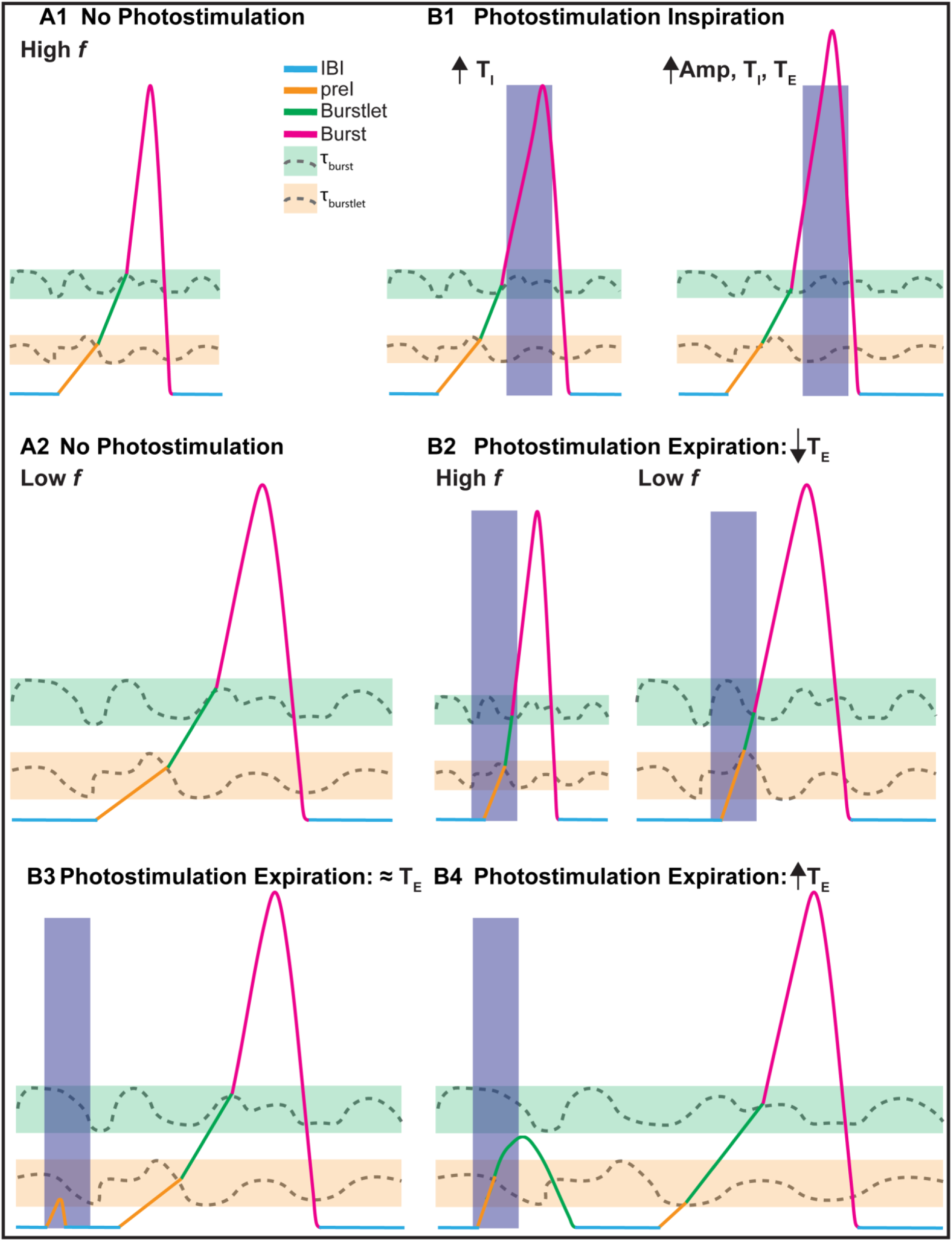
Proposed schema describing mechanisms underlying the phase- and state-dependent modulation of breathing by preBötC SST^+^ neurons (see text for details). Orange bar represents range of fluctuating thresholds (dotted line) for burstlet initiation, i.e., τ_burstlet_; green bar represents range of fluctuating thresholds (dotted line) for burst initiation, i.e., τ_burst_. Breathing cycles at **A1)** high *f* and **A2)** low *f* without photostimulation. Examples of breathing cycles with photostimulation (blue boxes) during **B1)** inspiration and **B2-4)** expiration. Each breathing cycle is subdivided into IBI (azure), preInspiration (orange), burstlet (green), and burst (red).

Notably, the effects of photostimulation during expiration transitioned from unimodal (phase advance) to bimodal (phase delay or phase advance) as *f* decreased below an apparent threshold of *f*≈2.5 Hz. Since these effects were seen when we normalized the variables in the test cycles by the preceding, unperturbed cycle, we suggest their *f*-dependency is a function of overall excitability rather than *f per se*.

The effects of photostimulation of preBötC SST^+^ neurons in anesthetized mice were phase- and state-dependent. This is consistent with optogenetic studies targeting preBötC SST^+^ neurons that showed phase-dependency (Cui *et al*., 2016) or other preBötC neurons that showed state-dependency (Doi and Ramirez, 2010; Montandon and Horner, 2013). We observed both phase- and state-dependency by examining the effects of phase-timed photostimulation in awake behaving mice breathing gas mixtures that were normal (∼room air: 21% O_2_, 0% CO_2_), hypoxic (8% O_2_, 0% CO_2_) or hypercapnic (21% O_2_, 5% CO_2_). While both hypoxia or hypercapnia increase respiratory network excitability, hypercapnia increases *f* (Guyenet and Bayliss, 2015; Del Negro, Funk and Feldman, 2018), whereas hypoxia decreases *f* (Del Negro, Funk and Feldman, 2018; Rajani *et al*., 2018). In all respiratory conditions without photostimulation (including normoxia) *f*≥2.5 Hz. With photostimulation in normoxia, hypoxia, or hypercapnia, in the first photostimulated breath following stimulus onset (typically ≤100 ms after stimulus onset), Amp increased, cycle duration decreased, and latency from pulse onset to inspiratory peak decreased, i.e., the phase advanced, a response similar to that seen in anesthetized mice when *f*≥2.5 Hz. Thus, photostimulation of preBötC SST^+^ neurons evoked a short latency excitatory effect regardless of experimental condition. The bimodal effect on T_E_ seen in anesthetized mice when baseline *f*<2.5 Hz was not observed in awake behaving mice, perhaps simply explained that in all cases baseline *f*≥2.5 Hz, regardless of whether *f* increased (hypercapnia) or *f* decreased (hypoxia) (see next paragraph). That low *f* was not seen in behaving mice could be explained by the absence of anesthesia, i.e., isoflurane, which affects neural excitability (Stuth, Stucke and Zuperku, 2012) to significantly lower *f*. We suggest that since in awake behaving mice the excitability state is higher than in anesthetized mice, the photostimulus was always sufficient to trigger a burst (Figure 10B2, left), i.e., advance the phase.

### Slower (>100 ms to 1 s) modulation

Prolonged (1 s) photostimulation in awake behaving mice, which spanned several breath cycles, decreased *f* compared to control (normoxia + 1 s pulse: 3.3 ± 0.2 Hz vs. normoxia: 4.4 ± 0.3 Hz, hypoxia + 1 s pulse: 2.0 ± 0.1 Hz, vs. hypoxia: 3.4 ± 0.1 Hz, hypercapnia + 1 s pulse: 3.8 ± 0.3 Hz vs. hypercapnia: 5.4 ± 0.2 Hz). Notably, of relevance to comparison with anesthetized mice, only prolonged photostimulation during hypoxia reduced *f*<2.5 Hz, while photostimulation during hypercapnia did not.

After the first photostimulated cycle, i.e., >100 ms to 1 s relative to pulse onset, responses were similar regardless of condition: i) no effect on the second photostimulated cycle; and ii) increased Amp on subsequent photostimulated cycles (Figure 8).

The mechanism underlying the absence of an effect on the second photostimulated breath is not obvious. We speculate that it could be an interaction between a short latency (≤100 ms) excitatory response and a slower (>100 ms) onset of an inhibitory response (perhaps by activation of the small subpopulation of inhibitory preBötC SST^+^ neurons), as proposed in anesthetized mice (Cui *et al*., 2016). Alternatively, increased Amp during the first photostimulated cycle might evoke a short-term synaptic depression, which affects the second photostimulated cycle, as similar mechanisms were described in Dbx1^+^ preBötC neurons (Kottick and Del Negro, 2015).

In the third and fourth photostimulated breaths during the 1 s stimulus, pattern (here measured as peak amplitude) was modulated while rhythm (here measured as the latency from pulse onset to inspiratory peak) was unaffected. Moreover, since responses in the awake states *f*<2.5 Hz (photostimulation during hypoxia) and *f*≥ 2.5 Hz (photostimulation during hypercapnia) were similar, but distinct under anesthesia, we suggest that state-dependency of SST-mediated modulation depends on overall breathing network excitability rather than *f*. That excitability state of *in vivo* experiments can affect breathing pattern and responses to perturbation could explain differences in experimental results, such as when comparing anesthetized vs. awake rodents, as is the case here, or when different type or level of anesthetics are used, e.g., mice anesthetized with a combination of ketamine, xylazine, and isoflurane (Cui *et al*., 2016) vs. isoflurane-anesthetized mice used here, or even when comparing results from anesthetized mice where peripheral oxygen sensors are denervated (Marchenko *et al*., 2016) vs. left intact (Janczewski *et al*., 2013).

In summary, activation of preBötC SST^+^ neurons differentially modulates breathing in a combinatory phase- and state-dependent manner and potentially via two distinct functional pathways as proposed by Cui et al (Cui *et al*., 2016). Here, a rapid SST-mediated pathway affects the first photostimulated breath and is excitatory, evoking a sigh-like or ectopic breath, and a slower SST-mediated pathway can be manifested as excitatory and/or inhibitory and modulates both Amp and *f*. Moreover, our results indicate that state-dependency is a function of overall excitability rather than simply on *f*. We surmise, therefore, that these two pathways are dependent on the state of the network excitability and are likely manifestations of intrinsic interactions among neuronal properties and synaptic and network dynamics.

While state and phase dependencies have been described independently, they were not previously considered as combinatory variables or as probabilistic. Here, we describe a biphasic effect of photostimulation throughout expiration that is probabilistic, i.e., photostimulation given at the same phase in consecutive cycles can give opposite response (lengthening vs. shortening of the phase) (Figure 5B1-B2). This novel data of probabilistic state- and phase-dependent responses to photostimulation presents properties of the preBötC that have not been predicted and cannot be readily accounted for in current models of preBötC pattern generation (Rybak *et al*., 2004; Shao, Tsao and Butera, 2006; Purvis *et al*., 2007; Smith *et al*., 2007; Jasinski *et al*., 2013).

This study *in vivo* in mice is an important extension in our understanding of generation of breathing pattern in mammals, in particular, the role of preBötC SST^+^ neurons. Moreover, the experimental protocols used reveal dynamic threshold-based mechanisms underlying rhythm generation in the context of the burstlet theory (Feldman and Kam, 2015; Ashhad and Feldman, 2020; Kallurkar *et al*., 2020). While partitioning our data at f≈2.5 Hz allowed us to differentiate two potentially distinct states of overall excitability, where the same stimulus produced distinct responses, we acknowledge that the relationship between baseline *f* and overall excitability in an intact behaving animal is complex and remains yet to be determined. We suggest that the burstlet/burst thresholds are higher and/or preBötC excitability is lower in lower excitability states, i.e., low *f*, therefore the probability that optogenetic manipulations result in sufficient network activity and synchronization to produce a burst, and, therefore, advance the phase, is lower.

What then is the role of preBötC SST^+^ neurons? Breathing is a robust and highly labile behavior that rapidly adjusts to constantly changing internal and external demands. preBötC SST^+^ neurons, in particular, have extensive reciprocal interconnectivity with multiple brains regions, e.g., brainstem bCPG and suprapontine nuclei (Tan *et al*., 2010; Yang and Feldman, 2018; Yang *et al*., 2020), which puts them in a key position to both process and transmit modulatory signals that affect breathing pattern. These observations underlie our conclusion that preBötC SST^+^ neurons are a significant contributor to the extraordinary and essential lability of breathing pattern and are likely involved in a broad range of breathing responses, such as in startle, exercise, stress, disease, and changes of altitude.

## Acknowledgments

We thank Carolina Thörn Perez and Sufyan Ashhad for useful comments and suggestions. This work was supported by National Institutes of Health grants NS07221 and HL135779.

## Supplementary Figures

**Figure S1.**
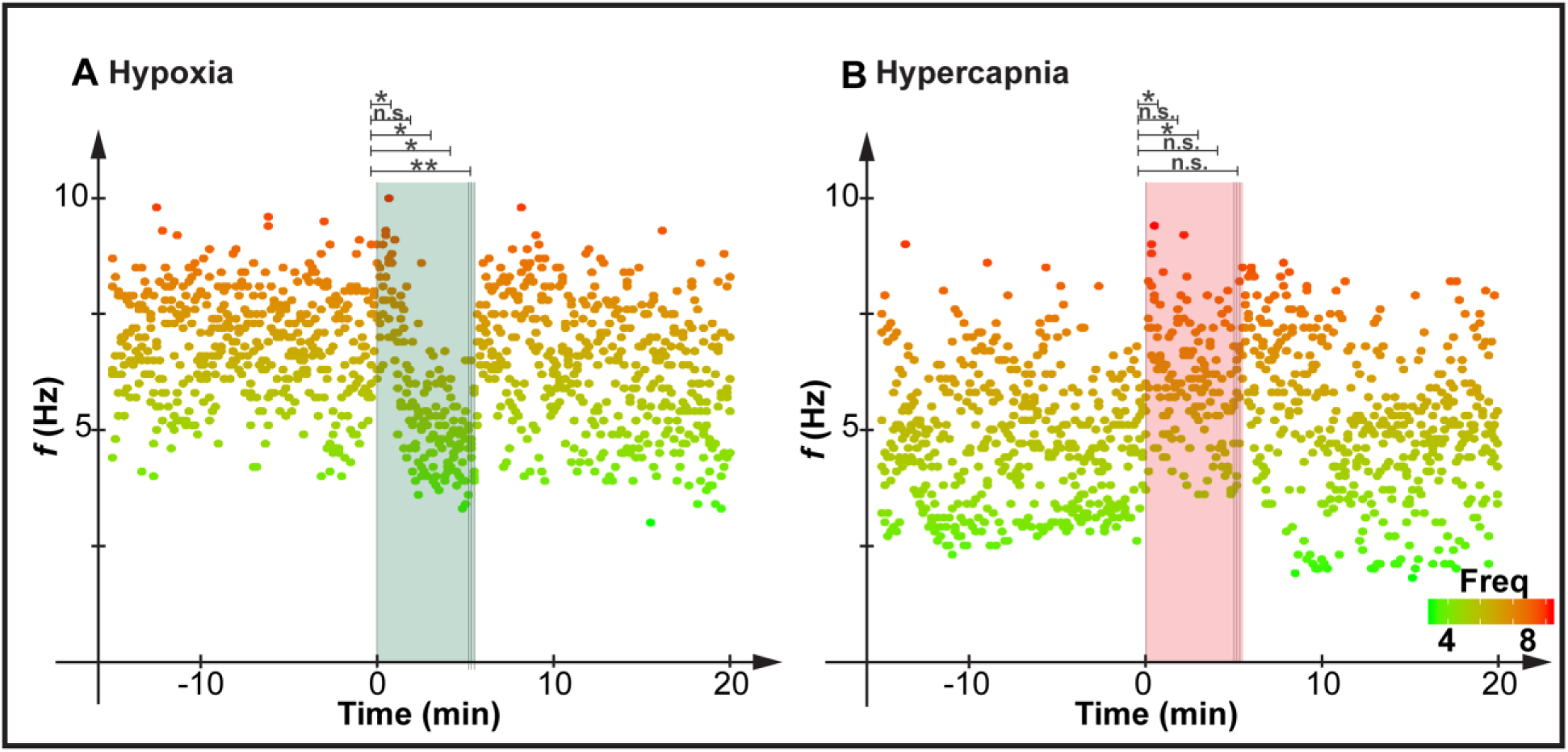
Control group without photostimulation. Mice were exposed to **A)** 5 mins hypoxia (5 mice) and **B)** 5 mins hypercapnia (5 mice). *F* (10 s bin) with a color code ranging from green (low *f*) to red (high *f*). **Right)** Post hoc Tukey’s multiple comparisons of 60 s bins; complete list of comparisons in Table S2B. ns if p≥0.05, * if 10^−5^≤p<0.05 and ** if p<10^−5^.

## Supplementary Tables

**Table S1.** Phase- and state-dependent modulation of T_I_, Amp, and T_E_ by preBötC SST^+^ neurons. For each phase bin, T_I_, Amp, and T_E_ (mean±SEM) for Pre, i.e., cycle prior to the photostimulated cycle, and Stim, i.e., photostimulated cycle when *f*<2.5Hz or *f* ≥2.5Hz. Unpaired t-tests: Pre vs. Stim. n.s. if p-value≥0.05.

| Phase | $T_i$ (ms) | | | | | |
| --- | --- | --- | --- | --- | --- | --- |
| | $f < 2.5$ Hz | | | $f \geq 2.5$ Hz | | |
| | Mean±SEM | | p-value | Mean±SEM | | p-value |
|  | Pre | Stim | Pre vs. Stim | Pre | Stim | Pre vs. Stim |
| $0 \leq \phi < 0.1$ | 163±2 | 168±2 | 0.03 | 134±2 | 147±2 | $10^{-5}$ |
| $0.1 \leq \phi < 0.2$ | 156±2 | 163±2 | 0.02 | 134±3 | 158±5 | $10^{-4}$ |
| $0.2 \leq \phi < 0.3$ | 153±2 | 161±3 | 0.03 | 137±4 | 171±8 | $10^{-4}$ |
| $0.3 \leq \phi < 0.4$ | 157±2 | 156±2 | n.s. | 133±2 | 133±4 | n.s. |
| $0.4 \leq \phi < 0.5$ | 155±2 | 153±2 | n.s. | 137±3 | 133±2 | n.s. |
| $0.5 \leq \phi < 0.6$ | 158±2 | 156±2 | n.s. | 131±2 | 130±2 | n.s. |
| $0.6 \leq \phi < 0.7$ | 155±2 | 154±2 | n.s. | 137±2 | 136±2 | n.s. |
| $0.7 \leq \phi < 0.8$ | 153±1 | 152±1 | n.s. | 133±2 | 132±2 | n.s. |
| $0.8 \leq \phi < 0.9$ | 159±2 | 160±2 | n.s. | 137±2 | 135±2 | n.s. |
| $0.9 \leq \phi < 1$ | 154±2 | 155±2 | n.s. | 140±2 | 139±2 | n.s. |
| **Phase** | **Amp (a.u.)** |  |  |  |  |  |
|  | **$f < 2.5$ Hz** |  |  | **$f \geq 2.5$ Hz** |  |  |
|  | Mean±SEM |  | p-value | Mean±SEM |  | p-value |
|  | Pre | Stim | Pre vs. Stim | Pre | Stim | Pre vs. Stim |
| $0 \leq \phi < 0.1$ | 33±1 | 41±2 | $10^{-3}$ | 27±2 | 38±2 | $10^{-5}$ |
| $0.1 \leq \phi < 0.2$ | 28±1 | 34±2 | 0.03 | 31±2 | 47±4 | $10^{-3}$ |
| $0.2 \leq \phi < 0.3$ | 31±2 | 34±2 | n.s. | 34±3 | 48±5 | 0.01 |
| $0.3 \leq \phi < 0.4$ | 28±1 | 27±1 | n.s. | 32±2 | 31±2 | n.s. |
| $0.4 \leq \phi < 0.5$ | 28±1 | 26±1 | n.s. | 34±2 | 31±1 | n.s. |
| $0.5 \leq \phi < 0.6$ | 26±2 | 25±1 | n.s. | 27±1 | 26±1 | n.s. |
| $0.6 \leq \phi < 0.7$ | 28±1 | 27±1 | n.s. | 30±1 | 29±1 | n.s. |
| $0.7 \leq \phi < 0.8$ | 29±1 | 28±1 | n.s. | 29±2 | 28±2 | n.s. |
| $0.8 \leq \phi < 0.9$ | 32±1 | 31±2 | n.s. | 30±1 | 28±1 | n.s. |
| $0.9 \leq \phi < 1$ | 30±2 | 30±2 | n.s. | 29±1 | 29±1 | n.s. |
| **Phase** | **$T_E$ (ms)** |  |  |  |  |  |
|  | **$f < 2.5$ Hz** |  |  | **$f \geq 2.5$ Hz** |  |  |
|  | Mean±SEM |  | p-value | Mean±SEM |  | p-value |
|  | Pre | Stim | Pre vs. Stim | Pre | Stim | Pre vs. Stim |
| $0 \leq \phi < 0.1$ | 513±17 | 532±18 | n.s. | 236±3 | 264±4 | $10^{-7}$ |
| $0.1 \leq \phi < 0.2$ | 586±23 | 620±24 | n.s. | 234±4 | 266±7 | $10^{-4}$ |
| $0.2 \leq \phi < 0.3$ | 504±21 | 485±24 | n.s. | 235±5 | 244±17 | n.s. |
| $0.3 \leq \phi < 0.4$ | 507±20 | 470±28 | n.s. | 244±3 | 138±15 | $10^{-7}$ |
| $0.4 \leq \phi < 0.5$ | 512±22 | 444±25 | n.s. | 235±4 | 115±12 | $10^{-10}$ |
| $0.5 \leq \phi < 0.6$ | 507±21 | 433±23 | 0.02 | 236±4 | 135±6 | $< 10^{-16}$ |
| $0.6 \leq \phi < 0.7$ | 527±21 | 462±22 | 0.04 | 238±3 | 149±4 | $< 10^{-16}$ |
| $0.7 \leq \phi < 0.8$ | 559±20 | 508±22 | n.s. | 241±4 | 184±4 | $10^{-15}$ |
| $0.8 \leq \phi < 0.9$ | 590±20 | 604±26 | n.s. | 238±4 | 208±4 | $10^{-7}$ |
| 0.9≤$\phi$<1 | 574±22 | 612±28 | n.s. | 234±4 | 224±4 | n.s. |

**Table S2A.** Breathing frequencies (mean±SEM; 1 minute bin) before (−15 to −1), during (0 to 4), and after (≥5) exposure to hypoxia or hypercapnia.

| Hypoxia |  |  |  | Hypercapnia |  |  |  |
| --- | --- | --- | --- | --- | --- | --- | --- |
| Time (min) | $f$ (Hz; mean±SEM) | Time (min) | $f$ (Hz; mean±SEM) | Time (min) | $f$ (Hz; mean±SEM) | Time (min) | $f$ (Hz; mean±SEM) |
| -15 | 6.7±0.2 | 0 | 8.0±0.2 | -15 | 4.9±0.3 | 0 | 6.6±0.3 |
| -14 | 7.0±0.2 | 1 | 6.4±0.2 | -14 | 5.2±0.2 | 1 | 5.8±0.2 |
| -13 | 6.6±0.2 | 2 | 5.2±0.2 | -13 | 4.2±0.2 | 2 | 6.3±0.2 |
| -12 | 6.9±0.2 | 3 | 5.1±0.2 | -12 | 4.4±0.3 | 3 | 5.9±0.2 |
| -11 | 6.8±0.2 | 4 | 4.7±0.1 | -11 | 4.6±0.3 | 4 | 5.5±0.2 |
| -10 | 6.9±0.2 | 5 | 5.4±0.2 | -10 | 5.0±0.3 | 5 | 6.4±0.2 |
| -9 | 7.1±0.2 | 6 | 7.0±0.2 | -9 | 5.1±0.3 | 6 | 6.0±0.3 |
| -8 | 7.1±0.2 | 7 | 6.8±0.2 | -8 | 4.8±0.2 | 7 | 6.1±0.2 |
| -7 | 6.8±0.2 | 8 | 7.1±0.3 | -7 | 4.9±0.2 | 8 | 5.8±0.3 |
| -6 | 6.9±0.2 | 9 | 6.8±0.3 | -6 | 5.2±0.3 | 9 | 5.5±0.4 |
| -5 | 6.7±0.1 | 10 | 6.6±0.2 | -5 | 4.7±0.3 | 10 | 4.9±0.3 |
| -4 | 7.1±0.2 | 11 | 6.3±0.2 | -4 | 4.3±0.2 | 11 | 4.7±0.2 |
| -3 | 6.5±0.3 | 12 | 6.4±0.3 | -3 | 4.5±0.2 | 12 | 4.5±0.2 |
| -2 | 6.4±0.2 | 13 | 6.2±0.2 | -2 | 4.6±0.2 | 13 | 4.8±0.3 |
| -1 | 6.7±0.2 | 14 | 6.1±0.2 | -1 | 4.8±0.2 | 14 | 4.4±0.2 |

**Table S2B.** Tukey’s Multiple Comparison Post Hoc Tests for all breathing frequencies between −15 and 15 minutes. Here, only shown comparisons between consecutive minutes. During hypoxia and hypercapnia, we compared the minute before hypoxia or hypercapnia (−1) and the following 5 minutes of exposure to the respiratory challenge (0 to 4).

| Before | Time (min) | Hypoxia Adj. p-value | Hypercapnia Adj. p-value | During | Time (min) | Hypoxia Adj. p-value | Hypercapnia Adj. p-value |
| --- | --- | --- | --- | --- | --- | --- | --- |
|  | -15 vs. -14 | n.s. | n.s. |  | -1 vs. 0 | 0.01 | 10<sup>-4</sup> |
|  | -14 vs. -13 | n.s. | n.s. |  | -1 vs. 1 | n.s. | n.s. |
|  | -13 vs. -12 | n.s. | n.s. |  | -1 vs. 2 | 10<sup>-4</sup> | 0.03 |
|  | -12 vs. -11 | n.s. | n.s. |  | -1 vs. 3 | 10<sup>-5</sup> | n.s. |
|  | -11 vs. -10 | n.s. | n.s. |  | -1 vs. 4 | 10<sup>-8</sup> | n.s. |
|  | -10 vs. -9 | n.s. | n.s. | **After** | 5 vs. 6 | 10<sup>-4</sup> | n.s. |
|  | -9 vs. -8 | n.s. | n.s. |  | 6 vs. 7 | n.s. | n.s. |
|  | -8 vs. -7 | n.s. | n.s. |  | 7 vs. 8 | n.s. | n.s. |
|  | -7 vs. -6 | n.s. | n.s. |  | 8 vs. 9 | n.s. | n.s. |
|  | -6 vs. -5 | n.s. | n.s. |  | 9 vs. 10 | n.s. | n.s. |
|  | -5 vs. -4 | n.s. | n.s. |  | 10 vs. 11 | n.s. | n.s. |
|  | -4 vs. -3 | n.s. | n.s. |  | 11 vs. 12 | n.s. | n.s. |
|  | -3 vs. -2 | n.s. | n.s. |  | 12 vs. 13 | n.s. | n.s. |
|  | -2 vs. -1 | n.s. | n.s. |  | 13 vs. 14 | n.s. | n.s. |
|  |  |  |  |  | 14 vs. 15 | n.s. | n.s. |

**Table S3A.** Breathing frequencies (Hz; mean±SE) of freely behaving mice in normoxia, hypoxia, and hypercapnia. Pre, Stim, and Post correspond to 1 s bins prior, during, and following photostimulation, respectively. We used the post-hoc Tukey’s test to perform multiple comparisons of breathing frequency among Pre, Stim, and Post (1 s bins) in normoxia, hypoxia, and hypercapnia. n.s. if p-value≥0.05.

|  | *f* (Hz; mean±SEM) |  |  |  | Adj. p-value |  |  |
| --- | --- | --- | --- | --- | --- | --- | --- |
|  | Normoxia | Hypoxia | Hypercapnia |  | Normoxia | Hypoxia | Hypercapnia |
| **Pre** | 4.4±0.3 | 3.4±0.1 | 5.4±0.2 | **Pre vs. Stim** | 10<sup>-3</sup> | 10<sup>-8</sup> | 10<sup>-5</sup> |
| **Stim** | 3.3±0.2 | 2.0±0.1 | 3.8±0.3 | **Pre vs. Post** | n.s. | n.s. | n.s. |
| **Post** | 5.1±0.3 | 3.5±0.2 | 5.9±0.2 | **Stim vs. Post** | 10<sup>-6</sup> | 10<sup>-9</sup> | 10<sup>-9</sup> |

**Table S3B.** We used the post-hoc Tukey’s test to perform multiple comparisons of breathing frequency among respiratory conditions (normoxia, hypoxia, and hypercapnia) during Pre, Stim, and Post. n.s. if p-value≥0.05.

| **Conditions** | Adj. p-value |  |  |
| --- | --- | --- | --- |
|  | Pre | Stim | Post |
| **Normoxia vs. Hypoxia** | 10<sup>-3</sup> | 10<sup>-5</sup> | 10<sup>-5</sup> |
| **Normoxia vs. Hypercapnia** | 10<sup>-3</sup> | n.s. | 0.05 |
| **Hypoxia vs. Hypercapnia** | 10<sup>-10</sup> | 10<sup>-9</sup> | 10<sup>-10</sup> |

**Table S4A.** Coefficients of Variation (CV) for Amp, Latency, and Duration (mean±SEM, %) in normoxia, hypoxia, hypercapnia. Adj. p-values calculated with Tukey’s Multiple Comparison Post Hoc Test. n.s. if p-value≥0.05.

|  | Condition | CV<sub>Amp</sub><br>(%; mean±SEM) | Conditions | Adj. p-value |
| --- | --- | --- | --- | --- |
| **Photostimulation** | Normoxia | 34±4 | Normoxia vs. Hypoxia | n.s. |
|  | Hypoxia | 42±8 | Normoxia vs. Hypercapnia | n.s. |
|  | Hypercapnia | 28±3 | Hypoxia vs. Hypercapnia | n.s. |
| **No Photostimulation** | Normoxia | 23±2 | Normoxia vs. Hypoxia | n.s. |
|  | Hypoxia | 16±4 | Normoxia vs. Hypercapnia | n.s. |
|  | Hypercapnia | 17±2 | Hypoxia vs. Hypercapnia | n.s. |
|  | Condition | CV<sub>Latency</sub><br>(%; mean±SEM) | Conditions | Adj. p-value |
| **Photostimulation** | Normoxia | 77±3 | Normoxia vs. Hypoxia | n.s. |
|  | Hypoxia | 82±4 | Normoxia vs. Hypercapnia | n.s. |
|  | Hypercapnia | 74±2 | Hypoxia vs. Hypercapnia | n.s. |
| **No Photostimulation** | Normoxia | 62±2 | Normoxia vs. Hypoxia | n.s. |
|  | Hypoxia | 70±3 | Normoxia vs. Hypercapnia | n.s. |
|  | Hypercapnia | 66±2 | Hypoxia vs. Hypercapnia | 0.02 |
|  | Condition | CV<sub>Duration</sub><br>(%; mean±SEM) | Conditions | Adj. p-value |
| **Photostimulation** | Normoxia | 41±6 | Normoxia vs. Hypoxia | n.s. |
|  | Hypoxia | 42±5 | Normoxia vs. Hypercapnia | n.s. |
|  | Hypercapnia | 40±4 | Hypoxia vs. Hypercapnia | n.s. |
| **No Photostimulation** | Normoxia | 29±4 | Normoxia vs. Hypoxia | 10<sup>-3</sup> |
|  | Hypoxia | 16±2 | Normoxia vs. Hypercapnia | 10<sup>-3</sup> |
|  | Hypercapnia | 16±2 | Hypoxia vs. Hypercapnia | n.s. |

**Table S4B.**
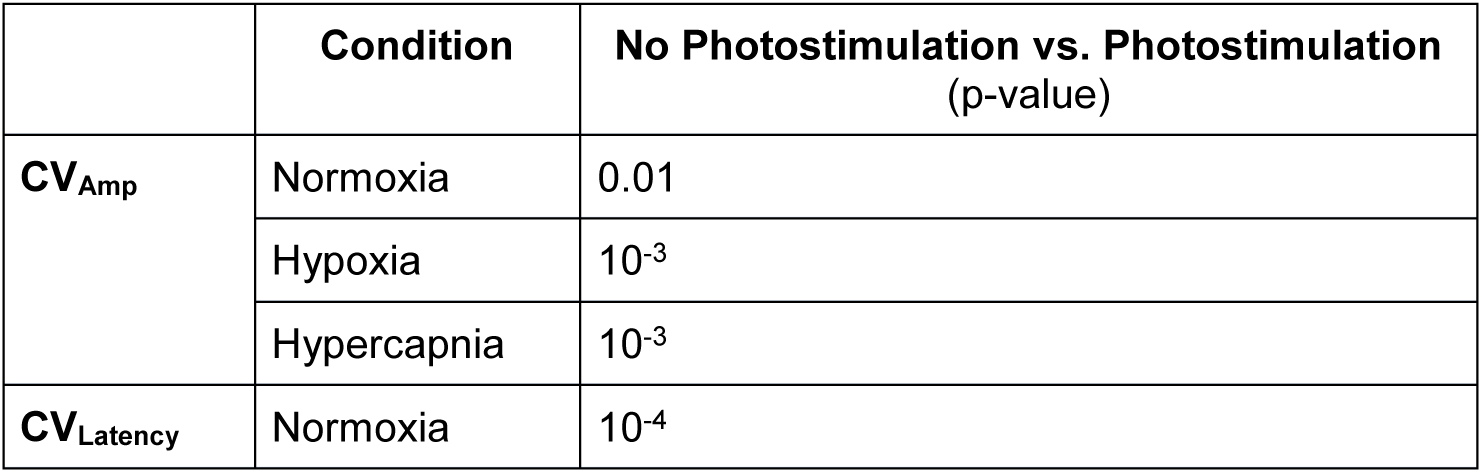
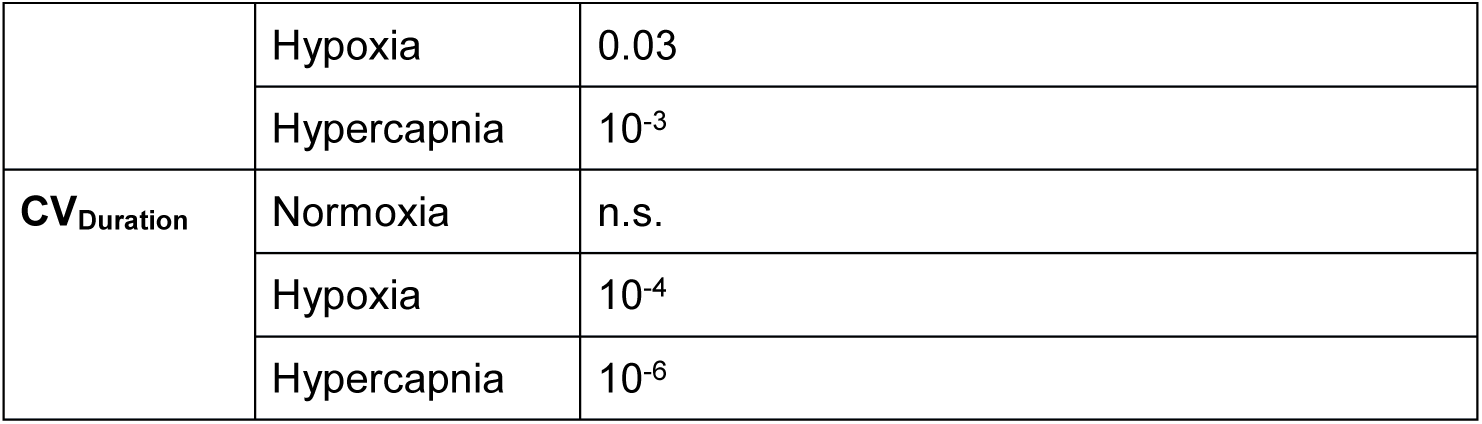
Unpaired t-tests photostimulation vs. no photostimulation: CV_Amp_, CV_Latency_, and CV_Duration_. n.s. if p-value≥0.05.

**Table S5A.**
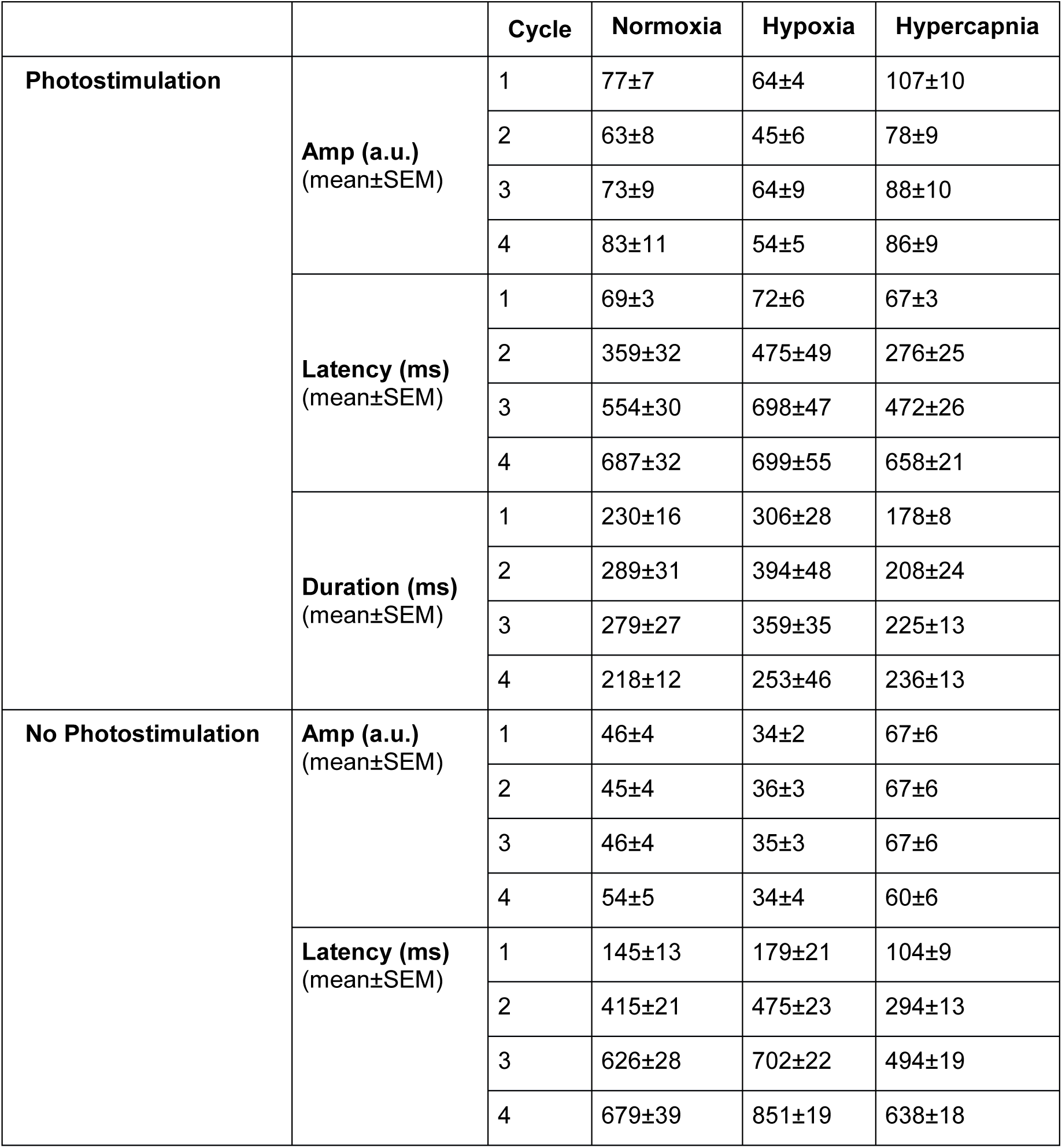
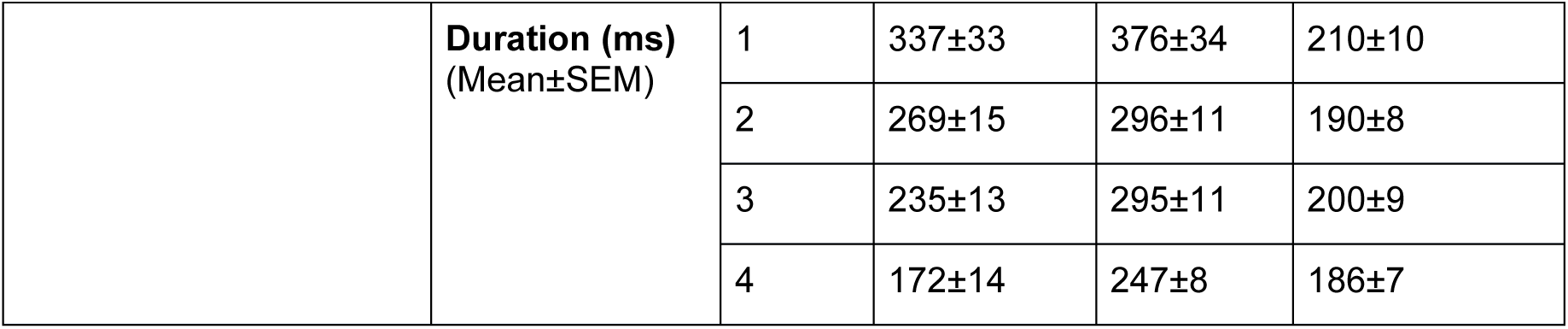
Amp, Latency, and Duration (mean±SEM) for each breathing cycle (1 to 4) in normoxia, hypoxia, and hypercapnia, with and without photostimulation.

|  |  | Cycle | Normoxia | Hypoxia | Hypercapnia |
| --- | --- | --- | --- | --- | --- |
| **Photostimulation** | **Amp (a.u.)**<br>(mean±SEM) | 1 | 77±7 | 64±4 | 107±10 |
|  |  | 2 | 63±8 | 45±6 | 78±9 |
|  |  | 3 | 73±9 | 64±9 | 88±10 |
|  |  | 4 | 83±11 | 54±5 | 86±9 |
|  | **Latency (ms)**<br>(mean±SEM) | 1 | 69±3 | 72±6 | 67±3 |
|  |  | 2 | 359±32 | 475±49 | 276±25 |
|  |  | 3 | 554±30 | 698±47 | 472±26 |
|  |  | 4 | 687±32 | 699±55 | 658±21 |
|  | **Duration (ms)**<br>(mean±SEM) | 1 | 230±16 | 306±28 | 178±8 |
|  |  | 2 | 289±31 | 394±48 | 208±24 |
|  |  | 3 | 279±27 | 359±35 | 225±13 |
|  |  | 4 | 218±12 | 253±46 | 236±13 |
| **No Photostimulation** | **Amp (a.u.)**<br>(mean±SEM) | 1 | 46±4 | 34±2 | 67±6 |
|  |  | 2 | 45±4 | 36±3 | 67±6 |
|  |  | 3 | 46±4 | 35±3 | 67±6 |
|  |  | 4 | 54±5 | 34±4 | 60±6 |
|  | **Latency (ms)**<br>(mean±SEM) | 1 | 145±13 | 179±21 | 104±9 |
|  |  | 2 | 415±21 | 475±23 | 294±13 |
|  |  | 3 | 626±28 | 702±22 | 494±19 |
|  |  | 4 | 679±39 | 851±19 | 638±18 |
|  | **Duration (ms)**<br>(Mean±SEM) | 1 | 337±33 | 376±34 | 210±10 |
|  |  | 2 | 269±15 | 296±11 | 190±8 |
|  |  | 3 | 235±13 | 295±11 | 200±9 |
|  |  | 4 | 172±14 | 247±8 | 186±7 |

**Table S5B.** Unpaired t-tests photostimulation vs. no photostimulation: Amp, Latency, and Duration for breathing cycles 1 to 4 in normoxia, hypoxia, hypercapnia. n.s. if p≥0.05.

|  | **Cycle** | **Normoxia** | **Hypoxia** | **Hypercapnia** |
| --- | --- | --- | --- | --- |
|  |  | No Photostimulation vs. Photostimulation (p-value) |  |  |
| **Amp** | 1 | 10<sup>-4</sup> | 10<sup>-8</sup> | 10<sup>-4</sup> |
|  | 2 | n.s. | n.s. | n.s. |
|  | 3 | 10<sup>-3</sup> | 10<sup>-3</sup> | n.s. |
|  | 4 | 0.02 | 0.01 | 0.02 |
| **Latency** | 1 | 10<sup>-7</sup> | 10<sup>-6</sup> | 10<sup>-5</sup> |
|  | 2 | n.s. | n.s. | n.s. |
|  | 3 | n.s. | n.s. | n.s. |
|  | 4 | n.s. | n.s. | n.s. |
| **Duration** | 1 | 10<sup>-3</sup> | n.s. | 0.02 |
|  | 2 | n.s. | n.s. | n.s. |
|  | 3 | n.s. | n.s. | n.s. |
|  | 4 | 0.01 | n.s. | 10<sup>-3</sup> |

